# Quantification and modeling of turnover dynamics of *de novo* transcripts in *Drosophila melanogaster*

**DOI:** 10.1101/2023.02.13.528330

**Authors:** Anna Grandchamp, Peter Czuppon, Erich Bornberg-Bauer

**Affiliations:** Institute for Evolution and Biodiversity, University of Münster, Münster, Germany; Department of Protein Evolution, Max Planck Institute for Biology, Tübingen, Germany

**Author notes:** these authors contributed equally to this work (Anna Grandchamp, Peter Czuppon).

## Abstract

Most of the transcribed eukaryotic genomes are composed of non-coding transcripts. Among these transcripts, some are newly transcribed when compared to outgroups and are referred to as *de novo* transcripts. *De novo* transcripts have been shown to play a major role in *de novo* gene emergence. However, little is known about the rates at which *de novo* transcripts are gained and lost in individuals of the same species. Here, we address this gap and estimate for the first time the *de novo* transcript turnover rate. We use DNA long reads and RNA short reads from seven samples of inbred individuals of *Drosophila melanogaster* to detect *de novo* transcripts that are (transiently) gained on a short evolutionary time scale. Overall, each sampled individual contains between 2,320 and 2,809 unspliced *de novo* transcripts with most of them being sample specific. We estimate that around 0.15 transcripts are gained per year, and that each gained transcript is lost at a rate around 5*×*10*^−^*^5^ per year. This high turnover of transcripts suggests frequent exploration of new genomic sequences within species. These rates provide first empirical estimates to better predict and comprehend the process of *de novo* gene birth.

## Introduction

In most multicellular organisms, only a small fraction of the genome codes for proteins (Sana et al., 2012; Piovesan et al., 2019). Intriguingly though, a large fraction of the non-genic genome is transcribed too, at least occasionally, e.g. under specific conditions such as stress or during development (Papantonis and Cook, 2010; Kim and Jinks-Robertson, 2012; van Steensel and Furlong, 2019). This production of transcripts throughout the entire genome has been described as pervasive transcription (Clark et al., 2011; Hangauer et al., 2013; Kellis et al., 2014) and has been demonstrated through several techniques (reviewed in Wade and Grainger, 2014). Many non-genic transcripts perform important functions. For example, rRNA and tRNA are indispensable constituents for the protein assembly by the ribosome (Palazzo and Lee, 2015). Some long non-coding RNAs (transcripts with more than 200 nucleotides) are involved in the regulation of splicing (Romero-Barrios et al., 2018) and transcription in general (Mercer et al., 2009; Ponting et al., 2009; Wang and Chang, 2011; Wang et al., 2011). Among non-coding transcripts, some are found only in a single species or population without a known ortholog and can therefore be defined as *de novo* transcripts.

Interestingly, many *de novo* transcripts were find to bind to ribosomes, indicating that part of their sequence might contain *Open Reading Frames* (ORFs) that are likely translated into small proteins (Aspden et al., 2014; Bazzini et al., 2014; Ruiz-Orera et al., 2014; Couso and Patraquim, 2017; Zhang et al., 2019; Patraquim et al., 2022). This protein-coding potential of *de novo* transcripts makes them important precursors of *de novo* genes, which are coding genes that arise from previously non-coding genomic regions (Carvunis et al., 2012). Together with the gain of an *Open Reading Frame*, the gain of transcription is a fundamental feature of *de novo* gene emergence (Reinhardt et al., 2013; Schlötterer, 2015; Van Oss and Carvunis, 2019; Schmitz and Bornberg-Bauer, 2017; Zhao et al., 2014). To understand the evolution and assess the rate of *de novo* gene emergence, it is therefore critical to quantify the evolutionary dynamics of *de novo* transcripts (Albà and Castresana, 2007; Schmitz and Bornberg-Bauer, 2017; Schmitz et al., 2018; Heames et al., 2020).

How quickly *de novo* transcripts are gained and lost is still largely unknown. Studies across multiple species indicate high turnover, i.e. gain or loss, of long non-coding RNAs (reviewed in Kapusta and Feschotte, 2014). For example, Necsulea et al. (2014) estimate that around 10,000 *de novo* transcripts emerged and were fixed in primates since the split from rodents around 100 million years (MY) ago, indicating a fixation rate of 100 transcripts per MY. A similar analysis, based on a comparison of transcriptomes from several rodent species, found a fixation rate of 5-10 transcripts per MY (Kutter et al., 2012). In these studies, transcripts were compared between species separated by large phylogenetic distances. This provides important insight into the transcript turnover in conserved genomic regions. However, a large amount of transcripts could not be compared, because conserved genomic regions only cover a small fraction of the genome due to genome evolution and reshuffling. Moreover, the total turnover of transcripts is estimated without an underlying evolutionary model of transcript gain and loss. For example, the difference between the number of transcripts in rat and mouse genomes can be either due to a loss in the mouse species or a gain (and subsequent fixation) in the rat species, which is impossible to distinguish with the existing data (Kutter et al., 2012).

Here, we aim to overcome these limitations by using a tightly controlled setting and very short phylogenetic distances. We use deep sequencing data from seven genomes of individual samples from a single species: *Drosophila melanogaster*. These genomes were extracted from inbred lines from different geographic locations. Because of the close evolutionary relationship of these samples, genomic rearrangements are highly unlikely to impact alignments and syntenies. We assembled a genome and a transcriptome for each homozygote DNA and RNA of the inbred lines, and used comparative genomics approaches to detect newly emerged transcripts in each sample. We additionally determined the genomic location of the *de novo* transcripts and determined orthology with newly proposed transcript-specific orthogroup definitions. The occurrence of transcripts in different samples allows us to infer transient dynamics of transcripts, i.e. their gain and loss rates. We call the birth of a transcript a *gain*, which importantly does not imply its fixation in the population. It may well be, and is probably much more likely, that a newly gained transcript will be lost again before it is fixed in the population or species. Transient gain and loss rates were estimated using the infinitely many genes model (Baumdicker et al., 2010). In contrast to previous studies, these estimates of gain and loss account for the evolutionary turnover of transcripts between samples of a single species, instead of solely the fixation (or loss) between different species. In addition to these transient processes, we also estimate the number of transcripts that have become fixed in our sample, in comparison to close Dipteran outgroups. Our study therefore provides a detailed picture of transcript turnover rates and evolutionary dynamics, which is important information to understand the process of *de novo* gene birth.

## Methods

### Sample-specific reference genomes

RNA short reads from whole genome illumina sequencing and DNA long reads from whole genome nanopore sequencing of seven isofemale lines of *D. melanogaster* were downloaded from NCBI (accession PRJNA929424). Among the seven lines, six come from European countries: Finland (FI), Sweden (SE), Denmark (DK), Spain (ES), Ukraine (UA) and Turkey (TR), and one is from Zambia (ZI) (SI, Section A.1). The RNA-Seq samples were built based on two males, two females and one larvae pooled together (Grandchamp et al., 2022b). The DNA was extracted from 50 individuals per inbred line. For each inbred line, we refer to the genetic material that was extracted and assembled from the line by a *sample*. The sample from Zambia is considered as the ancestral outgroup as the *D. melanogaster* from sub-Saharan Africa diverged from the European populations around 12,843 years ago (Li and Stephan, 2006; Laurent et al., 2011).

Our model and pipeline was set up to assess transcript gain and loss within a species. Reference genomes were assembled for each sample by mapping the long DNA read of the sample to a reference genome of *Drosophila melanogaster*, and extracting the 7 consensus genomes. This methodology has two advantages. First, it allows to compare the precise location of transcripts between each sample, and indeed allows to use the genome annotation to compare transcription between samples. It also made possible for us to build three definitions of transcript orthology with more comparable genomic location than from *de novo* assemblies. Second, as most studies of population genomics only use one reference genome for a species, our method can directly be applied in this context. However, the choice of not using the *de novo* assembled genomes also has a cost. We suspect that part of the transcript assemblies have been lost (Assessed to 130 unspliced *de novo* transcripts, SI Sections A.4 and A.5). Sample-specific duplication events inside genomes are lost, and so are the putative corresponding transcripts. For each sample, DNA reads shorter than 100 base pairs (bp) were removed from the long DNA reads by using Fitlong (github rrwick/fitlong). We used the genome of *D. melanogaster* BDGP6.28, downloaded from Ensembl (Yates et al., 2020), as a reference genome. DNA long reads from each samples were mapped to the reference genome using BWA-MEM (Houtgast et al., 2018) and a consensus genome per sample was extracted. The resulting SAM files were converted into BAM with Samtools view (Danecek et al., 2021). The BAM files were sorted and indexed with Samtools-sort and Samtools-index. Mapping statistics were obtained with Samtools-flagstats (SI, Section A.2), and alignments were visualized with *Integrative Genome Viewer* (IGV) (Thorvaldsdóttir et al., 2013). This procedure mapped 95-98% of the DNA to the reference genome. For each sample, a consensus genome was extracted, by using Samtools-mpileup (Ramirez-Gonzalez et al., 2012), bcftools (Narasimhan et al., 2016) and its function vcfutils.pl (Afgan et al., 2016) (Supplemental Deposit). The genomic regions that were not covered by mapping were completed by the corresponding region of the reference genome of *D. melanogaster*. Percentages of polymorphisms were assessed between samples, as well as genomic GC contents (SI, Section A.3). The polymorphism between samples was systematically lower than the threshold of 2%, confirming that the DNA belongs to a single species. This approach allowed us to detect genomic polymorphisms between samples, and to increase the precision for mapping sample-specific transcripts to genomes.

### Transcriptome assembly

The RNA reads of each sample were cleaned from adaptors with Trimmomatic (Bolger et al., 2014) and trimmed with FastQC (de Sena Brandine and Smith, 2019). The seven reference genomes were indexed with HISAT2 (Kim et al., 2019), and RNA reads from each sample were mapped to their respective reference genome with HISAT2 using the spliced aware option. The resulting SAM files were converted to BAM format with Samtools. The BAM files were sorted and indexed. In each sample, the transcriptomes were assembled with StringTie (Pertea et al., 2015) (Supplemental Deposit). The corresponding GTF and GFF files were built with TransDecoder from Trinity (Grabherr et al., 2011). A GTF file based on the transcriptome assembly was created with GTF to FASTA from GffRead (Pertea and Pertea, 2020) (Supplemental Deposit). For each sample, a final file was generated including the genomic position of unspliced transcripts, transcription orientation, and the size of the unspliced transcripts. Transcript coverage was recorded as *Transcripts Per Million* (TPM). The GTF file of the reference *D. melanogaster* genome was retrieved to access the position of established genes and transposable elements, which were annotated in the seven new reference genomes.

We estimated if the use of genomes assembled by mapping resulted in loss of *de novo* transcripts compared to *de novo* assembly. The unmapped transcripts were retrieved in each sample with bedtools and converted into FASTA. The unmapped transcripts were used as a query for a nucleotide BLAST search against a database of insects annotated TEs from RepeatMasker (http://www.repeatmasker.org). Moreover, the transcriptome assembly generated with *de novo* genome assemblies were retrieved from (Grandchamp et al., 2022b), and used as a BLAST query against the transcriptomes generated in this manuscript. We determined the genomic positions of transcripts without hits, and the ones corresponding to normal gene expression were annotated with bedtools.

### Detection of *de novo* transcripts

The transcriptomes of six Drosophila species (*Drosophila simulans, Drosophila sechellia, Drosophila virilis, Drosophila ananassae, Drosophila yakuba, Drosophila erecta*) were downloaded from Ensembl metazoa (http://metazoa.ensembl.org/index.html). These transcriptomes included all known protein-coding transcripts extracted from male and female Drosophila, as well as predicted coding gene transcripts. We also downloaded the whole set of annotated non-coding RNAs referenced for these species. To weed out as many transcripts of older origin as possible, these transcriptomes were merged and used as a target for nucleotide BLAST search with all sample-specific transcripts as query. Nucleotide BLASTs were performed with a cutoff of *E* = 10*^−^*^2^, in the forward direction of the target transcripts. Transcripts with no hit were considered as preliminary *de novo* transcripts. Preliminary *de novo* transcripts were then used as query to BLASTn search against the transcriptome of seven other Dipteran species: *Aedes aegypti, Anopheles arabiensis, Culex quiquefasciatus, Lucilia cuprina, Musca domestica, Stomoxys calcitrans, Teleopsis dalmanni*, downloaded from Ensembl, with a cutoff of *E* = 10*^−^*^2^, in the forward direction of the target transcripts. The remaining *de novo* transcripts without a BLAST hit were then filtered by TPM value. Transcripts with a TPM *<* 0.5 were removed from the data set to exclude low frequency transcripts. The remaining transcripts were considered as *de novo* transcripts (Supplemental Deposit). The same analysis was performed again with higher thresholds of expression of 1 and 5 TPM (data in SI, Section A.7). All of the follow-up analyses in the main text were performed on the *de novo* transcripts detected with a threshold of 0.5 TPM (TPM 1 and 5 in Sections A.7 and B.4).

The genomic positions of unspliced transcripts were retrieved from the seven GTF files generated from the transcriptome assemblies, and the overlap with genomic components was assessed with bedtools (Quinlan and Hall, 2010) and a python code developed for this purpose. The transcripts were distributed in six genomic positions: a) “Overlapping with an intergenic region and a fragment of a gene in the identical direction”, referred to as “exon longer”, b) “Overlapping with an intergenic region only”, referred to as “intergenic”, c) “Overlapping with an intergenic region and a fragment of a gene but in the opposite direction”, referred to as “antisense”, d) “Overlapping with an intergenic region and a pseudogene”, referred to as “pseudogenic”, e) “Overlapping with an intergenic region and an annotated non-coding RNA”, referred to as “ncRNA”, and f) “Inside of an intron” referred as “intronic”.

### Orthogroups of *de novo* transcripts

*De novo* transcripts from the seven samples were searched for orthology relationships between them. Constructing orthogroups of transcripts is more complex than constructing orthogroups of protein-coding genes. Two protein-coding genes are commonly grouped together into an orthogroup if the sequence similarity and coverage of their encoded protein exceeds a certain threshold, which suggests a common origin and a homologous function of the encoded protein. On the other hand, two transcripts can overlap but have arisen from a completely different transcription process if their transcription initiation and termination sites do not coincide. Moreover, two transcripts can have similar initiation and termination sites, but are differently spliced in different samples, giving rise to spliced transcripts with low sequence homology. Indeed, depending on the scientific question, transcript homology should be defined differently.

To cover a large number of possible scenario, we established three different definitions of transcript orthology that are depicted in Fig. 1:

**Figure 1:**
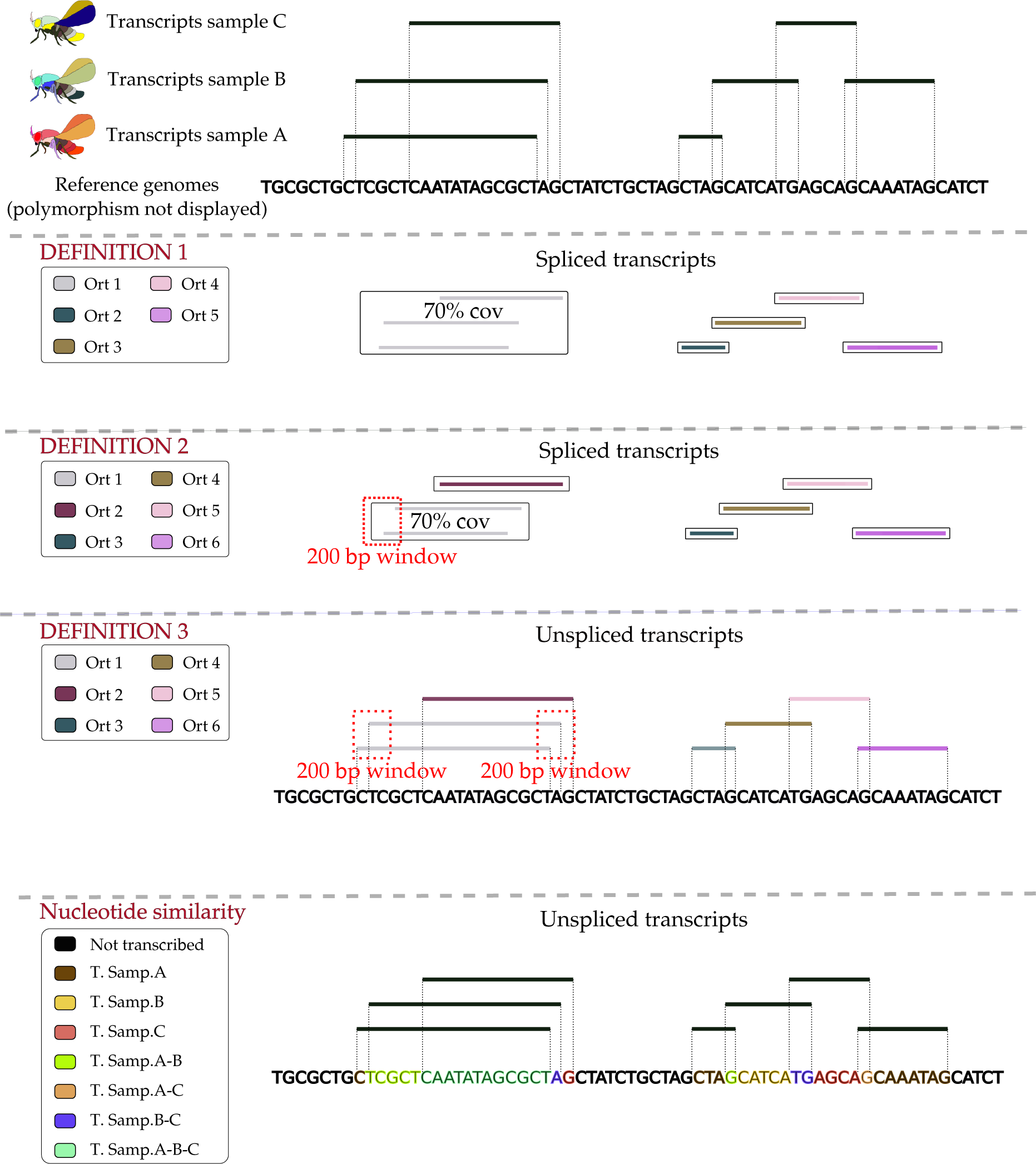
Transcript classifications. The figure illustrates the different *de novo* transcript definitions and the additional approach to categorize transcript nucleotides. In this example, only three samples are represented. The black nucleotides represent the reference genomes (all of the exact same size), the colored rows at the top show the transcripts.

> *Definition 1:* A set of transcripts are considered orthologous if their spliced sequences share at least 70% of coverage between all of the members of the orthogroup, and 75% of identity in reciprocal BLASTn.
>
> *Definition 2:* A set of transcripts are considered orthologous if their spliced sequences share at least 70% of coverage between all of the members of the orthogroup and 75% of identity in reciprocal BLASTn, and the transcription initiation sites of all of them can be found in a window of 500 bp of the genome.
>
> *Definition 3:* A set of transcripts are considered orthologous if their unspliced sequences start at a genomic position that is within a window of 500 bp, and end in a window of 500 bp.

*Definitions 1* and *2* compare spliced transcripts. *Definition 2*, in addition to sequence similarity, takes into account the genomic position of the transcription initiation site, thus making orthogroup affiliation more restrictive than *Definition 1*. In contrast, *Definition 3* builds on the initiation and termination sites but does not consider sequence similarity. In this definition, two splice variants of a single transcript would be clustered together, independently of the sequence similarity of the spliced variants. Indeed, if two transcripts emerge at the same position in two samples but with different splicing variants, they would still be detected by this definition.

To build the orthogroups following *Definitions 1* and *2*, *de novo* transcripts from each sample were used as target for BLASTn (Altschul et al., 1990) searches against the *de novo* transcripts of the six other samples. With a 70% coverage threshold, we did not encounter any ambiguous orthogroup definitions, e.g. when multiple transcripts overlapped with 70%, but not for all pairwise comparisons. A python script was used to sort the orthogroups according to our definitions. For *Definition 3*, a script was built to directly assess the overlapping positions and nucleotides of *de novo* transcripts between samples. To test the accuracy of our definitions, we used a set of 11,000 proto-genes from Grandchamp et al. (2022a), and distributed them into orthogroups by using the software OrthoFinder (Emms and Kelly, 2019) and our script for *Definition 1*. We found 93% of similarity between the results from OrthoFinder and our script (Orthofinder: 5687 orthogroups, Our script: 6124 orthogroups) (Supplemental Deposit). The difference of 7% is due to threshold of coverage between all members of an orthogroup implemented in our pipeline compared to OrthoFinder.

Additionally, we also used an alternative approach to estimate the similarity of transcripts between the samples. Instead of comparing transcripts with each other and treating them as a single object, we compare transcribed nucleotides. For example, if two transcripts of 200 bp found in two samples have an overlap of 50 bp, we will consider these 50 nucleotides as “*de novo* transcribed nucleotides” common to the two samples, and the remaining 150 nucleotides as “*de novo* transcribed nucleotides” specific to each sample. We used a python script to classify the nucleotides of transcripts into these categories. We refer to this approach as ‘Nucleotide similarity’.

### Evolutionary model of transient dynamics of *de novo* transcripts

We estimate gain and loss rates of *de novo* transcripts using a model that describes transcript gain and loss dynamics along the ancestry of the *D. melanogaster* samples. This evolutionary model is based on the *infinitely many genes model*, which has been developed to describe the gain and loss dynamics of genes in prokaryotes (Baumdicker et al., 2010, 2012). Analogously to the infinitely many genes model, in our model each specific *de novo* transcript arises only once during the short evolutionary time frame that we study. This means that any *de novo* transcript that is shared between different samples must have been gained in one of the ancestral branches common to both samples. Furthermore, in this model *de novo* transcripts are selectively neutral, i.e. they do not confer a fitness advantage or disadvantage, which is in line with empirical evidence (Palazzo and Koonin, 2020).

*De novo* transcripts are gained with probability *u* per unit of time and each transcript is lost with probability *v* per unit of time. Here, we use a generation as a unit of time. To translate between generations and years, we assume that the generation time of *D. melanogaster* is approximately two weeks (Fernández-Moreno et al., 2007). The total number of *de novo* transcripts in a sample after *t* units of time, denoted by *g*(*t*), is then described by the following dynamics (e.g. Eqs. (6), (7) in Collins and Higgs, 2012):

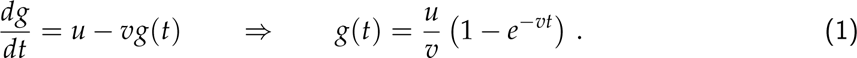

The equilibrium number of *de novo* transcripts therefore is *u*/*v*.

### Estimating *de novo* transcript gain and loss rates

Next, we outline how to estimate the gain and loss rates of *de novo* transcripts. To this end, we compare the empirical transcript frequency spectrum to a theoretical prediction of the frequency spectrum. The transcript frequency spectrum contains information about the number of transcripts shared by a certain number of samples. We denote by 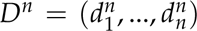 the empirical transcript frequency spectrum and by 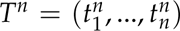 the theoretical prediction for the transcript frequency spectrum, where *n* is the number of samples that are studied. In the main text we restrict the study to the European samples, i.e. *n* = 6, whereas in Section B.3 in the SI we also include the Zambian sample in the analysis, i.e. *n* = 7. To estimate the parameters, we use a *χ*^2^ statistic to compare the empirical and theoretical frequency spectra, as has been done before (Baumdicker et al., 2012; Collins and Higgs, 2012):

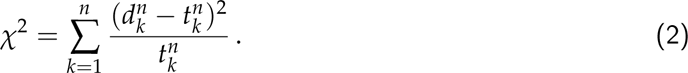

We now outline how to compute the theoretical transcript frequency spectrum following Baumdicker et al. (2010, 2012). The genealogy of the samples is modeled by a standard coalescent. The coalescent describes the ancestral relationship of samples taken from a neutrally evolving population of individuals in an unstructured population (Kingman, 1982); for a comprehensive introduction to coalescent theory we refer to Wakeley (2008). In this setting, the genealogy of the sample is given by a standard coalescent and the theoretical frequency spectrum can be computed analytically (Baumdicker et al., 2010). The European (meta-)population of *D. melanogaster* seems to have only a relatively weak population structure (Kapun et al., 2020), so that the genealogy of the European samples can be reasonably well described by a standard coalescent. We therefore restrict our analysis in the main text to these samples. The extended data set including the Zambian sample is analyzed in Section B.3 in the SI and shows no substantial difference in the estimated rates per generation (Table B.9 in the SI). Additionally, to study the robustness of the parameter estimation using coalescent theory, we also explore a model where the ancestry is given by the estimated phylogeny of the sample (Collins and Higgs, 2012) (SI, Section B.1). Using this alternative approach to estimate gain and loss rates, we do not find substantial differences to the reported estimates in Table 3 (details in the SI, Sections B.1 and B.3).

The frequency spectrum is given in gain and loss rates per generation. To transform these estimates to the time scale of the coalescent, we write *θ* = 2*N_e_u* and *ρ* = 2*N_e_v*, where *N_e_* denotes the effective population size. In our specific setting, *N_e_* corresponds to the effective population size of the European *D. melanogaster* population. Then *θ* is the average number of gained transcripts in 2*N_e_* generations and *ρ* is the rate at which a transcript is lost in 2*N_e_* generations. To transform these parameters to the scale of years, the estimated values are multiplied with the factor: *no. of generations per year* /(2*× effective population size*).

We denote by 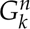 the number of transcripts shared by *k* samples out of the *n* samples. Similar to Collins and Higgs (2012), we also study two transcript classes: transcripts with high and low turnover. To distinguish their respective rates, we write *θ_s_*, *ρ_s_* for the “slow” class and *θ _f_* , *ρ _f_* for the “fast” class of transcripts. We drop the indices if we use only one class of transcripts in the following. The expected frequency spectrum, denoted by 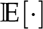, in its most general form is then given by (Baumdicker et al., 2010, Theorem 5 extended to multiple classes)

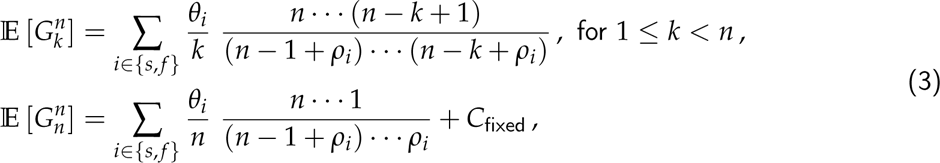

where *C*_fixed_ denotes the number of *de novo* transcripts that are fixed in our sample, which means that they have a loss rate equal to zero and are found in all samples.

### Numerical implementation of the parameter estimation

We used python and the integrated ‘minimize’ function from SciPy specifying the method ‘SLSQP’ to obtain parameter estimates through Eq. (2). We estimated parameters using one or two classes of transcripts plus the number of fixed transcripts, i.e. we estimated either three or five parameters. We constrained the parameters so that the mean number of transcripts per sample, the value *u*/*v* or *θ*/*ρ*, fits the empirical observation 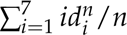. We did this to improve convergence of the minimization procedure by reducing the number of parameters to be estimated, which was necessary in the multi-class case. In addition, we conditioned the loss parameter(s) to be between 0 and 1,000 and the gain parameter(s) to be between 0 and 20,000.

Initial values for the parameters in the optimization routine were chosen by fitting the mean number of transcripts per sample and the pairwise-sample differences if we fitted one class of transcripts, as reasoned in Baumdicker et al. (2012) (more details in the SI, Section B.2). When fitting two classes of transcripts we divided the initial guess of the gain rate by two and set the initial gain and loss rate estimates of the slow transcript class to the initial values of the fast class divided by 100. All choices of initial parameter values are stated in Section B.2 in the SI.

### Programming and analyses

All statistical analyses were performed with R (R Core Team, 2022). Data reshuffling, analyses, orthology searches and modeling were performed with python (Van Rossum and Drake, 2009), and can be accessed at: https://github.com/AnnaGrBio/Transcripts-gain-and-loss.

## Results

### Sample-specific genome and transcriptome assemblies

The seven samples were extracted from seven inbred isolines of *Drosophila melanogaster* from different geographic locations (six from Europe, one from Zambia; details are provided in the Methods and Supplementary Information (SI), Section A.1). To this end, we compiled sample-specific reference genomes and identified *de novo* transcripts.

We mapped long DNA reads of each sample to the reference genome of *D. melanogaster* and extracted the seven consensus genomes. The percentage of DNA reads that correctly mapped to the reference genome ranged from 94.3% to 97.42% (SI, Section A.2 and Supplemental Deposit: https://doi.org/10.5281/zenodo.7681079). The percentages of SNPs between the aligned genomes of the seven samples ranged from 0.22% to 0.58% (SI, Section A.3), showing very low divergence as expected between samples of the same species. The Zambian sample diverged most from the other samples, which is consistent with its geographic separation from the European samples.

### *De novo* transcripts in samples

RNA reads from each of the seven individual samples were then mapped to their respective reference genome to build sample-specific transcriptomes (Supplemental Deposit)(SI, Section A.2). *De novo* transcripts were identified in the seven samples of *D. melanogaster* (details in the Methods section). In total, an average of 28,021 transcripts were found per sample (Table 1). Among these, on average 2,901 transcripts per sample were identified as *de novo* transcripts, as they showed no detectable homology to transcripts in other Diptera species, and had a required minimal expression threshold of 0.5 *Transcripts Per Million* (TPM) (Table 1). Among these transcripts, some arose via alternative splicing from a single unspliced precursor. When merging splicing variants as a single transcript, the average number of *de novo* transcripts dropped by 14.5% to an average of 2,481 per sample (Table 1). *De novo* transcripts were also defined with a higher threshold of expression. When using 1 TPM as a minimum level of expression of *de novo* transcripts, on average 871 transcripts were removed per transcriptome (SI, Section A.6), amounting to a total of 2,030 transcripts per sample. Genome assembly by DNA mapping can introduce a bias while assembling transcriptomes, as events of sample-specific indels and TEs are missing. Putative bias due to TEs insertions and transcript losses compared to *de novo* genome assembly were assessed (SI, Section A.4; A.5). We estimated that an average of 130 spliced (60 unspliced) *de novo* transcripts were lost in the assembly.

**Table 1:** Number of identified transcripts and *de novo* transcripts in the analysed samples.

| samples | DK<br>(Denmark) | ES<br>(Spain) | FI<br>(Finland) | SE<br>(Sweden) | TR<br>(Turkey) | UA<br>(Ukraine) | ZI<br>(Zambia) |
| --- | --- | --- | --- | --- | --- | --- | --- |
| # transcripts | 29,675 | 27,901 | 28,212 | 27,022 | 27,357 | 27,786 | 28,198 |
| # *de novo* transcripts | 2,908 | 2,842 | 2,708 | 2,714 | 2,997 | 3,024 | 3,116 |
| # *de novo* unspliced transcripts | 2,344 | 2,417 | 2,327 | 2,320 | 2,529 | 2,620 | 2,809 |

*De novo* transcripts were distributed uniformly across the chromosomes where they emerged (SI, Section A.3) and were found in all chromosomes except the mitochondrial one. According to their overlap with annotated genomic elements, we defined six different genomic positions to characterize the transcripts (details in the Methods section). Interestingly, *de novo* transcript distribution followed a similar pattern in all seven samples (Table 2): Around 60% of *de novo* transcripts overlapped with annotated genes in antisense, around 20% of the transcripts were found to be entirely intergenic, and just very few transcripts (around 3%) emerged inside introns. Except for these intronic transcripts, transcripts from all other genomic positions also overlapped, at least partially, with intergenic regions, making the intergenic region a very important pool of putative emergence of transcription initiation and termination. In addition, 8 to 12 *de novo* transcripts were consistently found to overlap with annotated pseudogenes in each sample. These pseudogenic transcripts were exceptionally long (11,000 to 150,000 nucleotides, Supplemental Deposit). Transcripts were also observed to overlap with non-coding RNA (ncRNA), but with an initiation site upstream or a termination site downstream of it. *De novo transcripts also contained introns (SI, Section A.7)*

**Table 2:** Number of *de novo* transcripts per sample ordered by genomic region.

| Position | DK | ES | FI | SE | TR | UA | ZI |
| --- | --- | --- | --- | --- | --- | --- | --- |
| number of *de novo* transcripts |  |  |  |  |  |  |  |
| exon longer | 307 | 355 | 352 | 351 | 362 | 366 | 396 |
| intergenic | 577 | 571 | 537 | 415 | 664 | 731 | 707 |
| intronic | 63 | 79 | 69 | 59 | 61 | 106 | 67 |
| ncRNA | 108 | 111 | 140 | 115 | 131 | 112 | 210 |
| pseudogene | 11 | 12 | 8 | 10 | 13 | 10 | 12 |
| antisense | 1842 | 1714 | 1602 | 1764 | 1766 | 1699 | 1724 |

**Table 3:** Estimated *de novo* transcript gain and loss rates. The gain and loss rates are measured as rates per year, the parameter *C*_fixed_ estimates the number of fixed transcripts in the sample.

| model | gain | loss | gain (slow) | loss (slow) | $C_{\text{fixed}}$ |
| --- | --- | --- | --- | --- | --- |
| **Definition 2** |  |  |  |  |  |
| 1 transcript class | 0.13 | $5 \times 10^{-5}$ | – | – | 91 |
| 2 transcript classes | 0.15 | $2 \times 10^{-4}$ | 0.06 | $3.1 \times 10^{-5}$ | 50 |
| **Definition 3** |  |  |  |  |  |
| 1 transcript class | 0.17 | $6.3 \times 10^{-5}$ | – | – | 95 |
| 2 transcript classes | 0.28 | $2.6 \times 10^{-4}$ | 0.05 | $3 \times 10^{-5}$ | 52 |

### Orthogroups of *de novo* transcripts

The number of identified orthogroups differed between the definitions (Fig. 2). *Definition 1* gave the smallest number of orthogroups (9,945), *Definition 2* clustered transcripts into 11,305 orthogroups, *Definition 3* into 12,223 orthogroups. *Definition 1* is solely based on transcript similarity and coverage and, contrarily to *Definition 2*, does not take into account the initiation position. Indeed, the fact that *Definition 1* defines less orthogroups suggests that around 2,000 transcripts overlap but have a substantially different initiation site. *Definition 3* characterizes orthogroups based only on the initiation and termination position of the unspliced transcripts, without considering splicing, similarity or coverage. The fact that this definition gave the highest number of orthogroups, combined with results from *Definitions 1 and 2*, suggests that *de novo* transcripts diverge by their initiation and termination location even though their sequences overlap. In other words, transcripts tend to emerge in nearby genomic regions, which makes them overlap, but at different transcription initiation sites, as they diverge more in their initiation position than in their coverage. Interestingly, when comparing the number of transcripts shared by samples, the three definitions showed the same frequency spectra (Fig. 2). Most orthogroups contain only one *de novo* transcript found in a single sample. The numbers of transcripts shared between samples decreases with the number of samples. This decrease is also visible for each genomic position of transcripts. As *Definition 1* is the least restrictive, it clusters more transcripts from different samples together. Accordingly, the number of transcripts shared by 2-7 samples was higher in *Definition 1* than in *Definitions 2* and *3*. The relative amount of transcripts specific to a single sample represented 53% of the total amount of orthogroups with *Definition 1*, 66% with *Definition 2* and 72% with *Definition 3*. The results from the alternative approach, which counts the number of nucleotides shared between the different samples, cannot be compared quantitatively to the three definitions as transcripts are not considered *per se*. Still, the frequency spectrum shows the same pattern as found by the three definitions.

**Figure 2:**
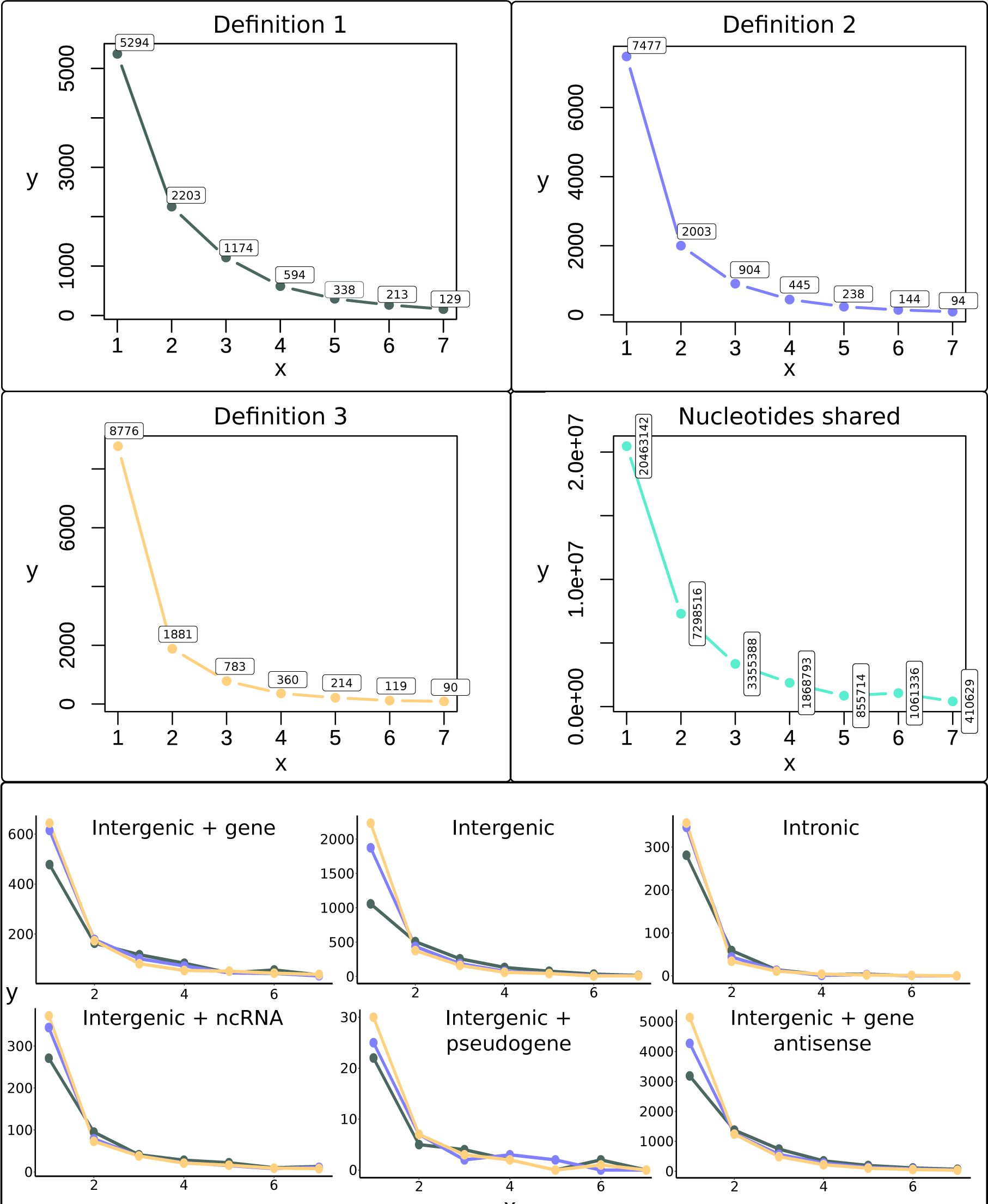
*De novo* transcripts shared by samples. The graphs from *Definitions 1, 2* and *3* show the number of transcripts shared by the number of samples they are found in. The graph called “Nucleotides shared” shows the number of nucleotides transcribed in common in one to seven sample. The *x*-axes correspond to the number of samples sharing a transcript, the *y*-axes show the number of transcripts. The six graphs at the bottom show the number of transcripts per genomic position shared between the samples. Each color represents a different definition: *Definition 1* in gray, *Definition 2* in blue, *Definition 3* in yellow.

The same definitions were used to cluster *de novo* transcripts into orthogroups by using *de novo* transcripts whose expression level was larger than 1 TPM (SI, Section A.8). The number of orthogroups were expectedly lower but the trend was the same as observed with the expression threshold of 0.5 TPM.

### Gain and loss rates of *de novo* transcripts

We estimated the gain and loss rates for the transcript frequency spectra obtained by *Definitions 2 and 3* (Fig. 2). The parameter estimation relies on comparison of a theoretically calculated transcript frequency spectrum and the empirically observed one (details in the Methods section). The estimated parameters are summarized in Table 3.

We explored two different models of transcript classes. In the simpler model, we consider a single class of transcripts, i.e. all transcripts are gained and lost at the same rates. The more complex model distinguishes between two classes of transcripts, a class with a high turnover rate, i.e. fast gain and model loss, and a class with a (relatively) lower turnover rate, i.e. slow gain and loss. Overall, the two transcript class model fits the observed frequency distribution slightly better, but the differences are small (Fig. 3). We therefore only discuss the estimates from the one transcript class model.

**Figure 3:**
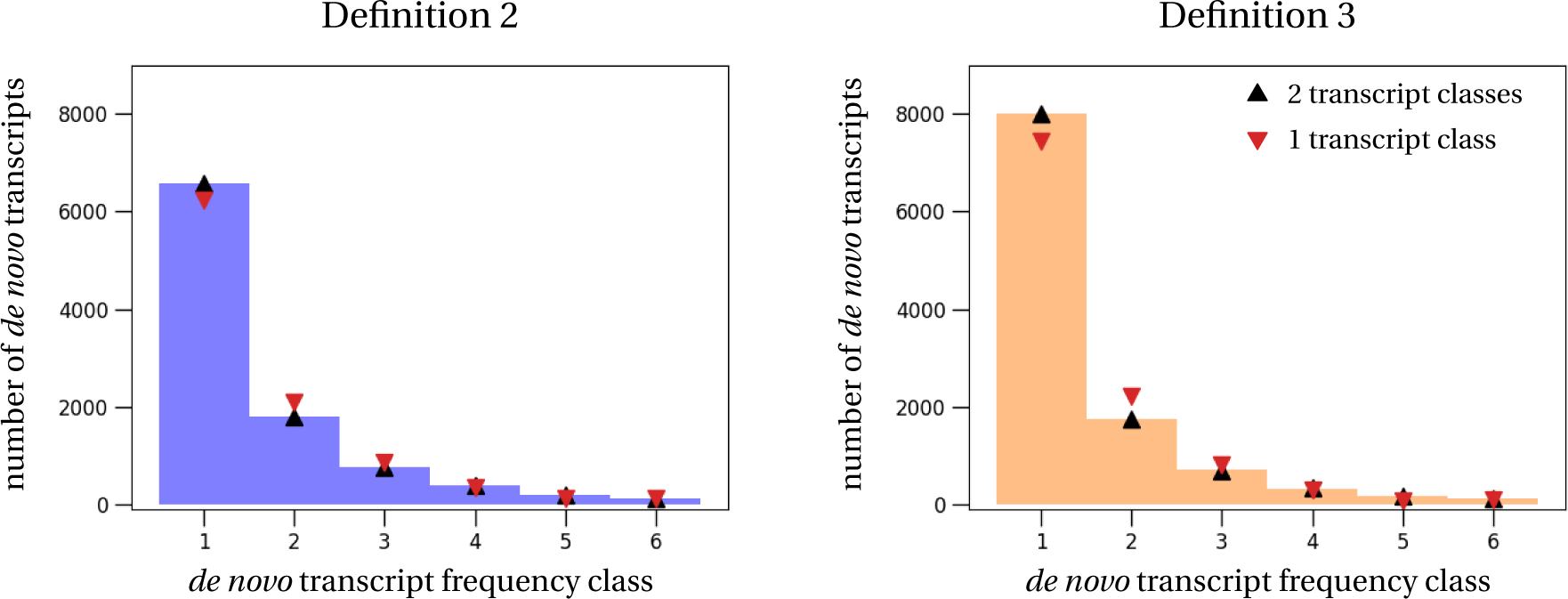
Empirical and estimated transcript frequency spectra. The histograms show the empirical *de novo* transcript frequencies obtained with *Definition 2* (left) and *Definition 3* (right) from the European samples only. Symbols show the theoretical frequency spectrum computed with the parameters estimated by the different models: black triangle – 2 transcript classes; red triangle (upside down) – 1 transcript class.

We find high gain and loss rates of transcripts, indicating high transient turnover dynamics of *de novo* transcripts. We estimate that between 0.13 and 0.17 new transcripts are gained per year, depending on the definition of transcript orthology and the number of transcript classes. Every single transcript is lost at a rate between 5 *×* 10*^−^*^5^ and 6.3 *×* 10*^−^*^5^ per year (fast and slow transcript classes), i.e. the expected life span of a transcript is approximately 20,000 years. We note that the gain and loss rates are estimated in rates per generation and need to be transformed to the per year scale (details in the Methods). This parameter transformation strongly depends on the assumptions of the generation time, here assumed to be two weeks (Fernández-Moreno et al., 2007), and the effective population size of the European *D. melanogaster* population, here set to 900,000 (Laurent et al., 2011). Uncertainty in these parameters can strongly impact the per year rate estimates in Table 3.

Lastly, we also estimate the number of transcripts that have become fixed in our data set. By fixed transcripts, we describe transcripts with the following properties: i) They are found in all samples, which means that they have been gained in the most recent common ancestor of the samples, and ii) they are modeled with a loss rate equal to zero. Adding this type of transcripts aids the parameter estimation by adding a degree of freedom to better describe the transcripts shared by all samples. We estimate 91-95 fixed transcripts in the one-transcript-class model and 50-52 fixed transcripts in the two-transcript-classes model. We emphasize that the number of fixed transcripts is sample-dependent. Even more, these transcripts are not necessarily fixed in the species, but may simply be found in all of the samples. Increasing the number of samples will likely reduce the number of estimated fixed transcripts because a newly added sample always has a positive probability of loosing one of the transcripts that is shared by all the other samples. For example, by including the Zambian sample, which is an outgroup to the European samples, into the data set, the estimated number of fixed transcripts reduces to 71-75, compared to the estimated 91-95 fixed transcripts within the European samples (Table B.9 in the SI).

### Transcript gain and loss rates per genomic region

We estimate the gain and loss rates separately for different genomic regions, using the same methodology as outlined in the previous section. Table 4 shows the estimated parameters (rates per year) for the one transcript class model. We find that the loss rates of transcripts are estimated between 2.6 *×* 10*^−^*^5^ and 5.5 *×* 10*^−^*^5^ across all genomic regions, which is consistent with the estimate from the aggregated data (Table 3). The gain rates, in contrast, differ more strongly over the genomic regions. One reason for this large variation might be that the genomic regions do not cover equal proportions of the genome. For example, intergenic regions cover around 68% of the *D. melanogaster* genome, while we approximate that the regions where transcripts overlap with non-coding RNAs only cover around 9% of the genome (details about the estimation of proportions are provided in the SI, Section B.5). Larger genomic coverage should therefore also result in a larger gain rate estimate because there are more positions where a *de novo* transcript could arise. To compare the gain rates across genomic regions in a meaningful way, we therefore normalize them according to their respective coverage. Strikingly, we find that the normalized gain rate of antisense transcripts is almost ten times larger than for all the other regions. Transcript gain in intronic regions is the lowest.

**Table 4:** Estimated *de novo* transcript gain and loss rates per genomic region. We have used the one transcript class model to estimate the gain and loss rates of transcripts and the fixed number of transcripts per genomic region. Parameter estimates of gain and loss rates are per year. To compare the gain rates between different genomic regions in a meaningful way, we have normalized the gain rates, denoted by gain_norm_, according to the coverage of the region in the genome. Normalization is done by rescaling the gain rates by their respective estimated coverage (details in SI, Section B.5).

| genomic region | Definition 2 |  |  |  | Definition 3 |  |  |  |
| --- | --- | --- | --- | --- | --- | --- | --- | --- |
|  | gain | gain<sub>norm</sub> | loss | C<sub>fixed</sub> | gain | gain<sub>norm</sub> | loss | C<sub>fixed</sub> |
| exon longer | $7.7 \times 10^{-3}$ | 0.05 | $2.6 \times 10^{-5}$ | 24 | 0.01 | 0.09 | $3.2 \times 10^{-5}$ | 31 |
| intergenic | 0.02 | 0.03 | $3.9 \times 10^{-5}$ | 1 | 0.03 | 0.05 | $4.4 \times 10^{-5}$ | 3 |
| intronic | $3.9 \times 10^{-3}$ | 0.02 | $5.5 \times 10^{-5}$ | 0 | $8.8 \times 10^{-3}$ | 0.04 | $4.5 \times 10^{-5}$ | 2 |
| ncRNA | $3 \times 10^{-4}$ | 0.04 | $3.6 \times 10^{-5}$ | 6 | $9.1 \times 10^{-3}$ | 0.1 | $3.8 \times 10^{-5}$ | 8 |
| antisense | 0.05 | 0.35 | $3.4 \times 10^{-5}$ | 2 | 0.07 | 0.45 | $4 \times 10^{-5}$ | 6 |

The estimated parameters from the alternative models and data sets are stated in SI, Table B.12. The general pattern remains the same, i.e. we find consistent loss rates across regions, and the lowest gain rate in intronic and the highest gain rate in antisense regions.

## Discussion

### De novo transcripts in Drosophila melanogaster

We investigated the emergence of *de novo* transcripts by using a unique setup based on samples of *Drosophila melanogaster* from different geographic locations. In each sample, between 2,708 and 3,116 transcripts with an expression level higher than 0.5 TPM showed no homology to any annotated transcript in Diptera and outgroup species, suggesting their *de novo* emergence (1,921 to 2,156 with expression level higher than 1 TPM). Some of these detected *de novo* transcripts are the result of alternative splicing, reducing the amount of unspliced *de novo* transcripts to 2,327–2,809 transcripts per sample. In total, their cumulative length covers 4–6% of the genome. We find that the gain of a transient *de novo* transcript is a frequent event. Previous studies already detected high amounts of new transcripts when comparing species or expression in different organs of the same species. For example, Brown et al. (2014) identified 1,875 new candidate long non-coding RNAs (lncRNAs) producing 3,085 transcripts in *D. melanogaster*, with 2,990 of them having no overlap with protein-coding genes of *D. melanogaster* or known lncRNAs in outgroup species. Huang et al. (2015) determined that 4.5 to 6.7% of transcripts detected in the transcriptome of *D. melanogaster* were un-annotated in flybase, which amounted to 1,669 transcripts derived from intronic regions and 2,192 from intergenic regions. We detected less *de novo* transcripts in *D. melanogaster* samples than these previous studies. However, the RNA of each of our samples included less developmental stages and full body transcripts compared to the other studies (tissues from larval, pupal and adults staged animals) . We particularly detected much fewer transcripts in intronic regions than the two previous studies, which could suggest that intronic *de novo* transcripts are specific to some developmental stages or tisues. Moreover, *de novo* genomes were assembled by mapping. Therefore, any transcript arising from a genome rearrangement may have been lost in our study. However, such an event remains unlikely as *de novo* genomes did not show major reshuffling event (Grandchamp et al., 2022b).

Strikingly, we find that in all samples most *de novo* transcripts (around 60%) overlap with coding genes in opposite direction. These antisense transcripts are a common phenomenon in genomes (Katayama et al., 2005). Their functions and mechanisms of emergence were reviewed in Barman et al. (2019). For example, antisense transcription plays an important role in gene expression regulation, as antisense transcripts can hybridize with forward transcripts and prevent translation of the forward-transcribed transcript (Pelechano and Steinmetz, 2013). Despite their importance in gene regulation, *de novo* emergence of antisense transcripts has not yet been intensively studied. Our results suggest that *de novo* emergence of antisense transcripts is common.

### High estimated rates of gain and loss suggest high transcript turnover

To understand and quantify the transient dynamics of *de novo* transcripts, we rely on the transcript frequency spectrum (Fig. 3), i.e. the number of transcripts shared between samples. The clustering of *de novo* transcripts into orthogroups forms the basis of this frequency spectrum. We distributed transcripts into orthogroups according to three new definitions that are based on different interpretations of orthology. All definitions resulted in similar patterns of transcripts shared across samples (Fig 2). Surprisingly to us, most *de novo* transcripts were specific to a single sample. This indicates a high transcript gain rate, which in turn suggests a high turnover of transcripts. We find this high rate of transcript gain independent of the specific TPM cutoff (main text: 0.5 TPM; SI, Section B.4: 1 TPM and 5 TPM). This result is of major importance because it suggests that initiation of transcription is easily gained within a species, but at different genomic locations in individuals. This observation should be taken into account when comparing transcript gains and losses between species, as the choice of the transcriptome representative of the species will impact the results of the comparison.

We quantified the transient dynamics, i.e. gain and loss rates, of *de novo* transcripts based on sequencing data from seven samples of *D. melanogaster*. We emphasize that most of the transcripts in our data are not fixed in *D. melanogaster*, i.e. we do not find them in all samples. Rather, the presence and absence pattern of transcripts throughout the different samples allows us to study the transient process where transcripts are on their way to fixation or extinction within the species. The term *gain* therefore refers to the emergence of a new transcript, but not to its fixation; similarly the term *loss* refers to the loss of a transcript within a sample, but not necessarily to its overall extinction on the species level. Additionally, we estimate the *number of fixed transcripts* in our sample, which are found in all samples and have an assumed loss rate equal to zero. To quantify the transient gain and loss processes, we used the infinitely many genes model (Baumdicker et al., 2010), and adapted it to the notion of transcripts. Independent of the definition, we find a high turnover rate of transcripts, i.e. high gain and loss rates. Using the data from *Definitions 2* and *3*, approximately 0.15 transcripts are gained per year, which is a factor of 10 smaller than a recent theoretical prediction based on a neutral evolutionary model (Iyengar and Bornberg-Bauer, 2023). A transcript is lost at a rate of *∼* 10*^−^*^4^ per year, which is consistent across all TPM threshold data sets. We find fewer orthogroups with *Definition 2* than with *Definition 3*, suggesting that new transcripts diverge rather in their initiation and termination sites than in their coverage. The high turnover of transcripts could be influenced by mutations in transcription factor motifs, strengthening or decreasing their ability to bind to the transcription machinery. That would be consistent with findings in previous studies (Ward and Kellis, 2012; Wade and Grainger, 2014; Young et al., 2015). For example, high turnover of initiation sites for transcription has been observed between human and mice (Frith et al., 2006), together with high reshuffling in upstream regulator regions (Brown and Feder, 2005; Odom et al., 2007; Wittkopp and Kalay, 2012; Cotney et al., 2013; Ballester et al., 2014; Vierstra et al., 2014; Villar et al., 2014, 2015). In *Bacillus subtilis*, 174 transcripts were found to have a new termination signal in the genome, further down than the original termination site, which had been inactivated (Nicolas et al., 2012). In fact, several studies in yeast and other eukaryotes demonstrated that modifications in transcription termination were involved in the abundant production of non-genic transcripts (Grosso et al., 2015; Rutkowski et al., 2015; Vilborg et al., 2015). To maintain constant transcript numbers over time, the large gain rate of transcription is countered by different transcript removal mechanisms. These mechanisms can be complementary and occur at different levels (Wade and Grainger, 2014; Lasa et al., 2011; Singh et al., 2014). For example, *de novo* transcripts can be directly degraded in the nucleus or the cytoplasm, as it has been shown in *Saccharomyces cerevisiae* (Jacquier, 2009; Porrua and Libri, 2015; Candelli et al., 2018b). In addition, transposable elements alter the landscape of pervasive transcription by pausing or terminating neighbouring transcription (Candelli et al., 2018a).

### Rates of *de novo* transcript gain vary across genomic regions

We classified *de novo* transcripts according to their genomic position. The proportion of transcripts found per genomic region was similar for all samples (Table 2). The loss rate of *de novo* transcripts was consistent across all genomic regions, which suggests that random mutations are driving the loss of transcription. This is further corroborated by the similar loss rate found in the data set with a 1 TPM threshold. In contrast, we found large differences between the gain rates in the different genomic regions (Table 4). The normalized gain rate was highest for *de novo* transcripts overlapping with genes in the opposite direction of their transcription (antisense). This finding is in line with studies showing that antisense transcription is frequent and is a major driver of evolution as it regulates genes expression (Katayama et al., 2005; He et al., 2008). Moreover, antisense transcripts can also be involved in diseases (Barman et al., 2019), and their frequent gain and loss could be adaptive in samples which were collected from different geographic locations, and thus possibly different environments. We also consistently find that the gain rate in intronic regions is lowest. Gain of a new intronic transcript might require both the acquisition of an initiation and a termination site inside an intron, contrarily to the other transcript types, which can potentially exploit an already existing site at one of the two ends. Additionally, the gain of these two elements has to occur in a region limited in size (the intron), which probably explains why gain of transcription inside an intron is less expected.

More unexpectedly, the gain of transcripts in intergenic regions and of transcripts overlapping with a gene occurs at a similar rate. We would have expected that transcripts emerging in overlap with a coding region have a higher chance to be quickly removed by purifying selection (Bourque et al., 2018). Transcripts overlapping with existing genes could, however, have a higher chance to acquire a coding function and by that become part of the gene splicing process, which might also explain the relatively high number of estimated fixed transcripts in this genomic region. These two counteracting processes, purifying selection vs. potentially beneficial coding function, could explain why the overall transcript gain rate in regions overlapping with genes in sense is comparable to the suspected selectively neutral dynamics in intergenic regions.

### Conclusion and future prospects

To summarize, we have estimated the transient dynamics of *de novo* transcripts in *D. melanogaster*. These estimates show that *de novo* transcripts are gained and lost at high rates inside this species. Gain rates vary across the genome, being highest in regions overlapping with genes in antisense and lowest in intronic regions. In contrast, loss rates of *de novo* transcripts are found to be similar across the genome.

Larger data sets will help to refine or confirm the generality of our findings. Including more samples, possibly from the same geographic regions or even the same generated isolines, would shed more light on the randomness and transience of *de novo* transcript gain and loss. Moreover, including genomes from several species that follow a phylogenetic gradient would enable modeling transcript dynamics beyond species boundaries. The time of divergence between species would provide insight into the fixation rates of transcripts in species under changing environmental conditions and after going through population bottlenecks.

In the broader context of genome evolution, our results are a first step to a more mechanistic and less phenomenological treatment and understanding of *de novo* transcript, and consequently *de novo* gene, evolutionary dynamics.

### Data access

The genomic DNA and RNA sequences are available under NCBI Bioproject PRJNA929424. Additionally, a package containing processed data is available in the zenodo archive: (https://doi.org/10.5281/zenodo.7681079), and is referred in the main text as “Supplemental Deposit”. The archive contains, for each sample, the polymorphic genomes, transcriptome assemblies, *de novo* transcripts, orthology comparison results, orthogroups according to the 4 classifications, proteins alignments for the phylogeny. Supplemental figures, informations, analyses and models are found in the Supplementary Information (SI). All programs are stored in the github link (https://github.com/AnnaGrBio/Transcripts-gain-and-loss).

## Competing interest statement

The authors declare no competing interests.

## Authors contribution

AG, PC and EBB conceptualised the study; AG assembled the genomes and transcriptomes, detected *de novo* transcripts, built the orthogroups and the phylogeny; PC conducted the mathematical modelling and parameter estimation; AG, PC and EBB validated and discussed the results; AG and PC wrote the original draft; AG, PC, EBB revised and commented on the draft; AG, PC and EBB acquired funding.

## Funding

AG and EBB acknowledge funding by Alexander von Humboldt-Stiftung. This work was supported in part by the Deutsche Forschungsgemeinschaft priority program “Genomic Basis of Evolutionary Innovations” (SPP 2349), project BO 2544/20-1 awarded to EBB and project CZ 294/1-1 awarded to PC.

## Notes

### Competing Interest Statement

The authors have declared no competing interest.

### Summary of Updates

The text has been reshuffled

