## Supplementary material for "Quantification and modeling of turnover dynamics of *de novo* transcripts in *Drosophila melanogaster*": SI

#### Contents

**A *De novo* transcripts** **S2**
- A.1 Populations origins . . . . . S2
- A.2 Mapping statistics . . . . . S2
  - A.2.1 DNA mapping statistics . . . . . S2
  - A.2.2 RNA mapping statistics . . . . . S4
- A.3 Genome comparison . . . . . S8
- A.4 percentage of unmapped RNA seq mapping to TEs . . . . . S8
- A.5 BLAST transcriptome of de novo genome against our transcriptome . . . . . S9
  - A.5.1 TPM 0.5 . . . . . S9
  - A.5.2 TPM 1 . . . . . S9
- A.6 Number of *de novo* transcripts per line with TPM of 1 as a cutoff . . . . . S10
- A.7 Introns in *de novo* transcripts . . . . . S10
- A.8 *de novo* transcripts orthogroups with TPM value of 1 TPM . . . . . S11
- A.9 Sample-specific phylogenetic trees . . . . . S14
  - A.9.1 BEAST tree . . . . . S14
  - A.9.2 RAxML tree . . . . . S15
  - A.9.3 Output from BEAST . . . . . S15

**B Details of the parameter estimation** **S18**
- B.1 Alternative method to estimate gain and loss rates . . . . . S18
- B.2 Initial parameter values for the parameter estimation . . . . . S19
- B.3 Results from all the estimation procedures . . . . . S19
- B.4 Results for alternative TPM threshold data sets . . . . . S19
- B.5 Normalization of gain rates per genomic region . . . . . S21
- B.6 Untransformed parameter estimates . . . . . S22

### A *De novo* transcripts

#### A.1 Populations origins

The lines used in the main text have been taken from seven populations, six from Europe and one from Zambia.

- DK: Karensminde, Denmark (DK; lat. 55.94, long. 10.21, alt. 16, September 2014, Mads Fristrup Schou)
- ES: Gimenells, Spain (ES; lat. 41.62, long. 0.62, alt. 173, September 2018, Josefa Gonzalez)
- FI: Akaa, Finland (FI; lat. 61.10, long. 23.52, alt. 110, July 2018, Maaria Kankare)
- SE: Lund, Sweden (SE; lat. 55.69, long. 13.19, alt. 28, August 2015, Jessica Abbott)
- TR: Yesiloz, Turkey (TR; lat. 40.23, long. 32.26, alt. 680, September 2018, Banu Önder)
- UA: Uman, Ukraine (UA; lat. 48.75, long. 30.21, alt. 214, August 2018, Iryna Kozeretska)
- ZI: Siavonga, Zambia (ZI, Line ZI418; lat. -16.32, long. 28.42, alt. 479, July 2010, John Pool)

#### A.2 Mapping statistics

Statistics from flagstag of DNA read mapping to the genomes and RNA reads mappings per lines.

##### *A.2.1 DNA mapping statistics*

- DK line

- 6271451 + 0 in total (QC-passed reads + QC-failed reads)
- 0 + 0 secondary
- 3004299 + 0 supplementary
- 0 + 0 duplicates
- 6102929 + 0 mapped (97.31%)
- 0 + 0 paired in sequencing
- 0 + 0 read1
- 0 + 0 read2
- 0 + 0 properly paired (N/A)
- 0 + 0 with itself and mate mapped
- 0 + 0 singletons (N/A)
- 0 + 0 with mate mapped to a different chr
- 0 + 0 with mate mapped to a different chr (mapQ>=5)

- ES line

- 6735241 + 0 in total (QC-passed reads + QC-failed reads)
- 0 + 0 secondary
- 2915171 + 0 supplementary
- 0 + 0 duplicates
- 6351539 + 0 mapped (94.30%)
- 0 + 0 paired in sequencing
- 0 + 0 read1

- 0 + 0 read2
- 0 + 0 properly paired (N/A)
- 0 + 0 with itself and mate mapped
- 0 + 0 singletons (N/A)
- 0 + 0 with mate mapped to a different chr
- 0 + 0 with mate mapped to a different chr (mapQ>=5)

• FI line

- 5494997 + 0 in total (QC-passed reads + QC-failed reads)
- 0 + 0 secondary
- 2691487 + 0 supplementary
- 0 + 0 duplicates
- 5353135 + 0 mapped (97.42%)
- 0 + 0 paired in sequencing
- 0 + 0 read1
- 0 + 0 read2
- 0 + 0 properly paired (N/A)
- 0 + 0 with itself and mate mapped
- 0 + 0 singletons (N/A)
- 0 + 0 with mate mapped to a different chr
- 0 + 0 with mate mapped to a different chr (mapQ>=5)

• SE line

- 12206908 + 0 in total (QC-passed reads + QC-failed reads)
- 0 + 0 secondary
- 3859789 + 0 supplementary
- 0 + 0 duplicates
- 11684378 + 0 mapped (95.72%)
- 0 + 0 paired in sequencing
- 0 + 0 read1
- 0 + 0 read2
- 0 + 0 properly paired (N/A)
- 0 + 0 with itself and mate mapped
- 0 + 0 singletons (N/A)
- 0 + 0 with mate mapped to a different chr
- 0 + 0 with mate mapped to a different chr (mapQ>=5)

• TR line

- 8186682 + 0 in total (QC-passed reads + QC-failed reads)
- 0 + 0 secondary
- 3975194 + 0 supplementary
- 0 + 0 duplicates
- 7828812 + 0 mapped (95.63%)
- 0 + 0 paired in sequencing
- 0 + 0 read1
- 0 + 0 read2
- 0 + 0 properly paired (N/A)

- 0 + 0 with itself and mate mapped
- 0 + 0 singletons (N/A)
- 0 + 0 with mate mapped to a different chr
- 0 + 0 with mate mapped to a different chr (mapQ>=5)

- UA line

- 4013604 + 0 in total (QC-passed reads + QC-failed reads)
- 0 + 0 secondary
- 2090231 + 0 supplementary
- 0 + 0 duplicates
- 3911816 + 0 mapped (97.46%)
- 0 + 0 paired in sequencing
- 0 + 0 read1
- 0 + 0 read2
- 0 + 0 properly paired (N/A)
- 0 + 0 with itself and mate mapped
- 0 + 0 singletons (N/A)
- 0 + 0 with mate mapped to a different chr
- 0 + 0 with mate mapped to a different chr (mapQ>=5)

- ZI line

- 4176096 + 0 in total (QC-passed reads + QC-failed reads)
- 0 + 0 secondary
- 1828575 + 0 supplementary
- 0 + 0 duplicates
- 3958524 + 0 mapped (94.79%)
- 0 + 0 paired in sequencing
- 0 + 0 read1
- 0 + 0 read2
- 0 + 0 properly paired (N/A)
- 0 + 0 with itself and mate mapped
- 0 + 0 singletons (N/A)
- 0 + 0 with mate mapped to a different chr
- 0 + 0 with mate mapped to a different chr (mapQ>=5)

##### *A.2.2 RNA mapping statistics*

- DK line

- 277384441 + 0 in total (QC-passed reads + QC-failed reads)
- 31033757 + 0 secondary
- 0 + 0 supplementary
- 0 + 0 duplicates
- 251494441 + 0 mapped (90.67%)
- 246350684 + 0 paired in sequencing
- 123175342 + 0 read1
- 123175342 + 0 read2
- 211595538 + 0 properly paired (85.89%)

- 216192696 + 0 with itself and mate mapped
- 4267988 + 0 singletons (1.73%)
- 478626 + 0 with mate mapped to a different chr
- 416152 + 0 with mate mapped to a different chr (mapQ>=5)

- ES line

- 235786625 + 0 in total (QC-passed reads + QC-failed reads)
- 26473645 + 0 secondary
- 0 + 0 supplementary
- 0 + 0 duplicates
- 209020460 + 0 mapped (88.65%)
- 209312980 + 0 paired in sequencing
- 104656490 + 0 read1
- 104656490 + 0 read2
- 175733272 + 0 properly paired (83.96%)
- 179239776 + 0 with itself and mate mapped
- 3307039 + 0 singletons (1.58%)
- 198914 + 0 with mate mapped to a different chr
- 167446 + 0 with mate mapped to a different chr (mapQ>=5)

- FI line

- 231847987 + 0 in total (QC-passed reads + QC-failed reads)
- 25560897 + 0 secondary
- 0 + 0 supplementary
- 0 + 0 duplicates
- 205768045 + 0 mapped (88.75%)
- 206287090 + 0 paired in sequencing
- 103143545 + 0 read1
- 103143545 + 0 read2
- 172581112 + 0 properly paired (83.66%)
- 176657538 + 0 with itself and mate mapped
- 3549610 + 0 singletons (1.72%)
- 405396 + 0 with mate mapped to a different chr
- 349456 + 0 with mate mapped to a different chr (mapQ>=5)

- SE line

- 241462405 + 0 in total (QC-passed reads + QC-failed reads)
- 34398567 + 0 secondary
- 0 + 0 supplementary
- 0 + 0 duplicates
- 226050184 + 0 mapped (93.62% )
- 207063838 + 0 paired in sequencing
- 103531919 + 0 read1
- 103531919 + 0 read2

- 184606354 + 0 properly paired (89.15%)
- 188461504 + 0 with itself and mate mapped
- 3190113 + 0 singletons (1.54%)
- 381308 + 0 with mate mapped to a different chr
- 332207 + 0 with mate mapped to a different chr (mapQ>=5)

• TR line

- 219529333 + 0 in total (QC-passed reads + QC-failed reads)
- 28239344 + 0 secondary
- 0 + 0 supplementary
- 0 + 0 duplicates
- 205110289 + 0 mapped (93.43%)
- 191289989 + 0 paired in sequencing
- 95644995 + 0 read1
- 95644994 + 0 read2
- 170334351 + 0 properly paired (89.05%)
- 173749805 + 0 with itself and mate mapped
- 3121140 + 0 singletons (1.63%)
- 226644 + 0 with mate mapped to a different chr
- 193137 + 0 with mate mapped to a different chr (mapQ>=5)

• UA line

- 224825641 + 0 in total (QC-passed reads + QC-failed reads)
- 28755635 + 0 secondary
- 0 + 0 supplementary
- 0 + 0 duplicates
- 207526297 + 0 mapped (92.31%)
- 196070006 + 0 paired in sequencing
- 98035003 + 0 read1
- 98035003 + 0 read2
- 171924686 + 0 properly paired (87.69%)
- 175467462 + 0 with itself and mate mapped
- 3303200 + 0 singletons (1.68%)
- 231726 + 0 with mate mapped to a different chr
- 196828 + 0 with mate mapped to a different chr (mapQ>=5)

• ZI line

- 188643800 + 0 in total (QC-passed reads + QC-failed reads)
- 28677593 + 0 secondary
- 0 + 0 supplementary
- 0 + 0 duplicates
- 175683086 + 0 mapped (93.13%)
- 159966207 + 0 paired in sequencing
- 79983104 + 0 read1

- 79983103 + 0 read2
- 141211755 + 0 properly paired (88.28%)
- 144479773 + 0 with itself and mate mapped
- 2525720 + 0 singletons (1.58%)
- 200496 + 0 with mate mapped to a different chr
- 169742 + 0 with mate mapped to a different chr (mapQ>=5)

#### A.3 Genome comparison

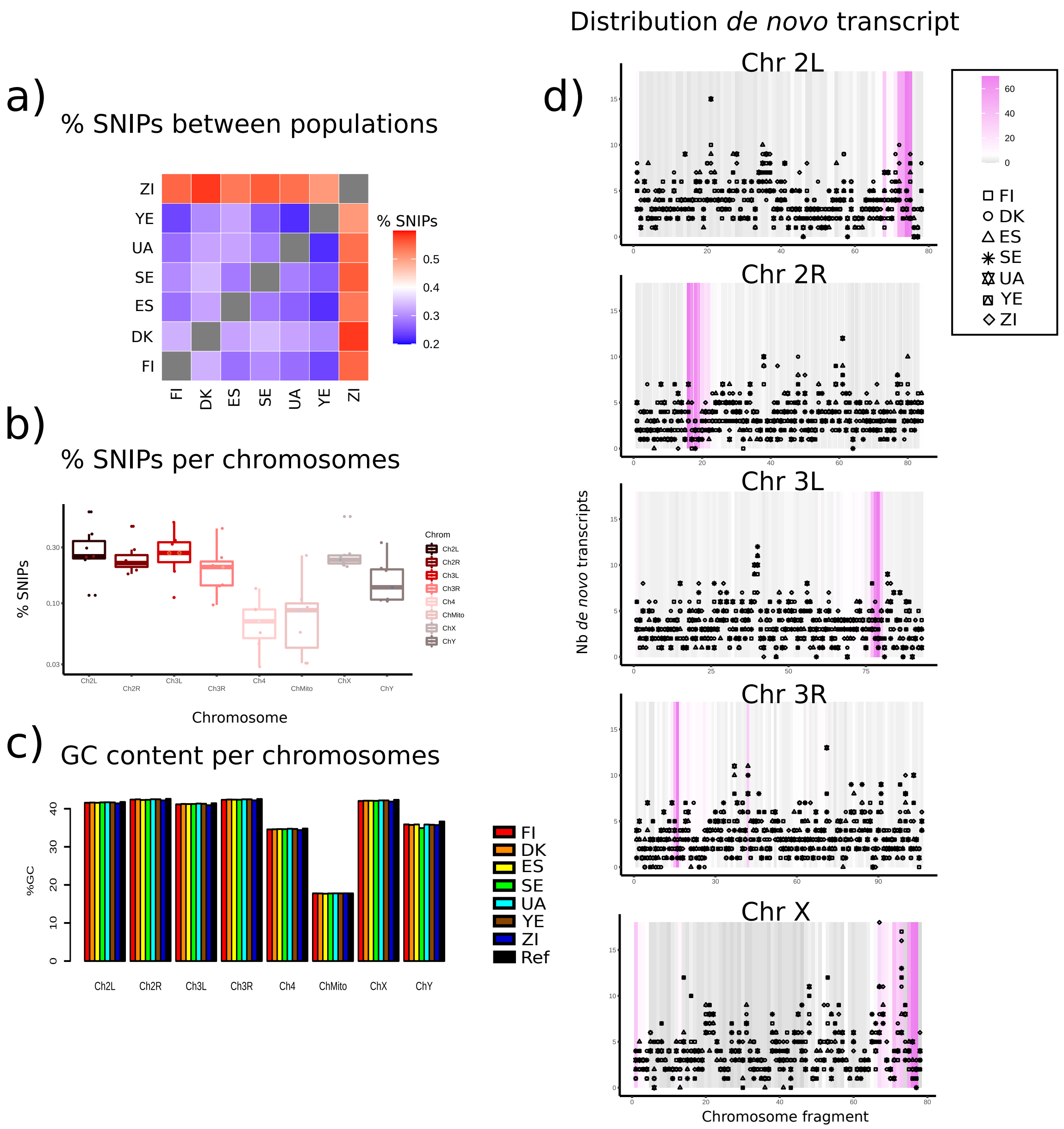


Figure S1: **Genome comparisons.** a) Mapping percentages, b) SNPs between genomes and SNPs between chromosomes, c) GC content, d) distribution of *de novo* transcripts in chromosomes.

#### A.4 percentage of unmapped RNA seq mapping to TEs

We assessed putative bias in the detection of *de novo* transcripts due to the method of assembly by mapping. DNA mapping covered an average of 99% of the genomes, indicating that the SNPs were accurately detected. The RNA mapping statistics were lower (88.75% to 93.62%) than for the RNA mapped to the corresponding genomes assembled *de novo*. We suspected that this difference could be explained by the multiplication of TEs inside samples. Unmapped transcripts were used as a BLAST target against fly TE databases, which showed that 10–18% of unmapped transcripts corresponded to

TEs: DK. 15.3%

ES. 9.4%

FI. 17.3%

SE. 13.9%

TR. 17%

UA. 15%

ZI. 16%

#### A.5 BLAST transcriptome of de novo genome against our transcriptome

We also assessed if the transcipts detected with the *de novo* assembly were correctly detected in our analyses (SI, Section A.5). Among the 30,000 transcripts detected on average in Grandchamp et al. (2022a), an average of 401 transcripts (470 spliced variants) could not be detected in our transcriptomes at an expression level larger than 0.5 (TPM), and 320 at an expression level larger than 1 TPM. Surprisingly, most of these transcripts were genic transcripts (average of 341 per samples), which suggests a putative loss of around 130 spliced *de novo* transcripts corresponding to 60 unspliced *de novo* transcripts.

##### *A.5.1 TPM 0.5*

FI total nb spliced transcripts: 494

FI total nb unspliced transcripts: 428

FI total nb spliced transcripts overlaping annotated element: 339

DK total nb spliced transcripts: 478

DK total nb unspliced transcripts: 384

DK total nb spliced transcripts overlaping annotated element: 330

ES total nb spliced transcripts: 460

ES total nb unspliced transcripts: 396

ES total nb spliced transcripts overlaping annotated element: 357

SE total nb spliced transcripts: 444

SE total nb unspliced transcripts: 374

SE total nb spliced transcripts overlaping annotated element: 335

UA total nb spliced transcripts: 541

UA total nb unspliced transcripts: 465

UA total nb spliced transcripts overlaping annotated element: 377

TR total nb spliced transcripts: 445

TR total nb unspliced transcripts: 386

TR total nb spliced transcripts overlaping annotated element: 342

ZI total nb spliced transcripts: 427

ZI total nb unspliced transcripts: 371

ZI total nb spliced transcripts overlaping annotated element: 309

##### *A.5.2 TPM 1*

FI total nb spliced transcripts: 391

FI total nb unspliced transcripts: 346

FI total nb spliced transcripts overlaping annotated element: 267

DK total nb spliced transcripts: 386

DK total nb unspliced transcripts: 316
DK total nb spliced transcripts overlaping annotated element: 262
ES total nb spliced transcripts: 363
ES total nb unspliced transcripts: 314
ES total nb spliced transcripts overlaping annotated element: 287
SE total nb spliced transcripts: 361
SE total nb unspliced transcripts: 305
SE total nb spliced transcripts overlaping annotated element: 273
UA total nb spliced transcripts: 411
UA total nb unspliced transcripts: 346
UA total nb spliced transcripts overlaping annotated element: 299
TR total nb spliced transcripts: 349
TR total nb unspliced transcripts: 314
TR total nb spliced transcripts overlaping annotated element: 279
ZI total nb spliced transcripts: 343
ZI total nb unspliced transcripts: 301
ZI total nb spliced transcripts overlaping annotated element: 254

#### A.6 Number of *de novo* transcripts per line with TPM of 1 as a cutoff

DK. 2,156
ES. 1,947
FI. 1,921
SE. 2,008
TR. 2,068
UA. 2,063
ZI. 2,050

#### A.7 Introns in *de novo* transcripts

Unexpectedly, splicing was observed in some *de novo* transcripts. We found 0.23 – 0.48 introns per *de novo* transcript on average, which is less than the average 1.8 introns per gene in the reference genome of *D. melanogaster* (Hiller et al., 2009). *De novo* transcripts overlapping with exons in forward sense of transcription (abbreviated by exon longer), or overlapping with genes but in the opposite direction of transcription (antisense), had the highest number of introns (average of 0.48 and 0.38, Table A.1). Intronic and intergenic transcripts had the lowest number of introns (average of 0.23 and 0.31). Still, the number of introns and of alternative splicing was on average low and smaller than the average number of introns from protein-coding genes. Other transcripts, like lncRNAs, are expected to become spliced by the same mechanisms as protein-coding transcripts (Krchňáková et al., 2019; Quinn and Chang, 2016). It is interesting to see that *de novo* transcripts are also capable of splicing, despite being newly emerged. Particularly, *de novo* transcripts that are antisense and overlapping with an existing gene in frame had the highest number of introns on average. Introns from the genes to which they overlap could thus putatively be reused by antisense transcripts. This would also explain why intronic and intergenic *de novo* transcripts had, in contrast, the lowest number of introns.

| position | DK | ES | FI | SE | TR | UA | ZI |
|---|---|---|---|---|---|---|---|
| **average number of introns** | | | | | | | |
| exon longer | 0.63 | 0.44 | 0.45 | 0.45 | 0.46 | 0.41 | 0.50 |
| intergenic | 0.35 | 0.36 | 0.26 | 0.33 | 0.30 | 0.32 | 0.26 |
| intronic | 0.27 | 0.22 | 0.17 | 0.22 | 0.20 | 0.19 | 0.36 |
| ncRNA | 0.44 | 0.27 | 0.27 | 0.34 | 0.32 | 0.29 | 0.35 |
| pseudogene | - | - | - | - | - | - | - |
| antisense | 0.42 | 0.37 | 0.38 | 0.36 | 0.41 | 0.38 | 0.34 |

Table A.1: **Number of introns per *de novo* transcripts.**

#### A.8 *de novo* transcripts orthogroups with TPM value of 1 TPM

| Total transcripts | Definition 1 | Definition 2 | Definition 3 |
|---|---|---|---|
| 1 line | 3,972 | 5,141 | 5,928 |
| 2 lines | 1,608 | 1,470 | 1,390 |
| 3 lines | 818 | 636 | 565 |
| 4 lines | 415 | 328 | 264 |
| 5 lines | 226 | 165 | 157 |
| 6 lines | 158 | 122 | 99 |
| 7 lines | 86 | 63 | 69 |

Table A.2: **Number of identified transcripts and *de novo* transcripts in the analysed lines with TPM ≥ 1**.

| ExonLonger | Definition 1 | Definition 2 | Definition 3 |
|---|---|---|---|
| 1 line | 374 | 481 | 498 |
| 2 lines | 154 | 158 | 153 |
| 3 lines | 93 | 82 | 67 |
| 4 lines | 68 | 64 | 46 |
| 5 lines | 41 | 39 | 49 |
| 6 lines | 51 | 41 | 38 |
| 7 lines | 31 | 27 | 34 |

Table A.3: **Number of identified transcripts and *de novo* transcripts overlapping with an exon in frame with a TPM ≥ 1**.

| Intergenic | Definition 1 | Definition 2 | Definition 3 |
|---|---|---|---|
| 1 line | 642 | 893 | 970 |
| 2 lines | 215 | 186 | 166 |
| 3 lines | 105 | 72 | 64 |
| 4 lines | 49 | 38 | 38 |
| 5 lines | 30 | 16 | 18 |
| 6 lines | 8 | 4 | 2 |
| 7 lines | 5 | 5 | 5 |

Table A.4: **Number of identified transcripts and *de novo* transcripts in an intergenic regions with a TPM ≥ 1**.

| Intronic | Definition 1 | Definition 2 | Definition 3 |
|---|---|---|---|
| 1 line | 139 | 161 | 156 |
| 2 lines | 22 | 17 | 18 |
| 3 lines | 5 | 6 | 5 |
| 4 lines | 2 | 1 | 3 |
| 5 lines | 0 | 0 | 0 |
| 6 lines | 0 | 0 | 0 |
| 7 lines | 0 | 0 | 0 |

Table A.5: **Number of identified transcripts and *de novo* transcripts inside an intron with a TPM ≥ 1**.

| ncRNA | Definition 1 | Definition 2 | Definition 3 |
|---|---|---|---|
| 1 line | 136 | 148 | 156 |
| 2 lines | 48 | 43 | 40 |
| 3 lines | 29 | 21 | 23 |
| 4 lines | 10 | 10 | 10 |
| 5 lines | 5 | 6 | 5 |
| 6 lines | 6 | 4 | 4 |
| 7 lines | 2 | 2 | 2 |

Table A.6: **Number of identified transcripts and *de novo* transcripts overlapping with an ncRNA with a TPM ≥ 1.**

| Pseudogene | Definition 1 | Definition 2 | Definition 3 |
|---|---|---|---|
| 1 line | 19 | 21 | 22 |
| 2 lines | 4 | 5 | 5 |
| 3 lines | 2 | 1 | 4 |
| 4 lines | 2 | 3 | 1 |
| 5 lines | 0 | 2 | 0 |
| 6 lines | 2 | 0 | 1 |
| 7 lines | 0 | 0 | 0 |

Table A.7: **Number of identified transcripts and *de novo* transcripts overlapping with a pseudogene with a TPM ≥ 1.**

| Antisense | Definition 1 | Definition 2 | Definition 3 |
|---|---|---|---|
| 1 line | 2,662 | 3,437 | 4,126 |
| 2 lines | 1,165 | 1,061 | 1,008 |
| 3 lines | 584 | 454 | 402 |
| 4 lines | 284 | 212 | 166 |
| 5 lines | 150 | 102 | 85 |
| 6 lines | 91 | 73 | 54 |
| 7 lines | 48 | 29 | 28 |

Table A.8: **Number of identified transcripts and *de novo* transcripts overlapping with an exon in antisense with a TPM ≥ 1.**

#### A.9 Sample-specific phylogenetic trees

Phylogenetic trees of the seven samples built with BEAST and RAXml.

##### *A.9.1 BEAST tree*

One of the estimation methods of *de novo* transcript gain and loss rates (explained in Section B.1) relies on a dated phylogenetic tree. To obtain an estimate of the phylogenetic history of the seven samples of *D. melanogaster*, we downloaded the genomes of each line assembled *de novo* in Grandchamp et al. (2022b). We did not use the reference genomes constructed by mapping in that study, as these genomes do not contain events of reshuffling, indels or transposable element insertions that are more precise to reconstruct the phylogenetic tree. Furthermore, we note that because of constant gene flow between populations, a strict phylogenetic classification in form of a binary tree is neither possible nor desirable but serves as a first approximation to group populations such that gain and loss rates can be estimated.

For each line, all coding sequences (CDS) were retrieved with GffRead (Pertea and Pertea, 2020). For all genes, longest CDS common to each line (11,568) were stored in orthogroups. Sequences from the 11,568 orthogroups were aligned with MAFFT (Katoh and Standley, 2013) and the 11,568 alignments were concatenated in one single alignment with python. This concatenated alignment was converted to nexus format with Seqmagick (fhcrc.github.io/seqmagick) and used as input for the software BEAST (Bouckaert et al., 2014; Drummond et al., 2012) for constructing a dated tree (details in SI, Section A.5).

To set the priors for constructing this tree, we used the software BEAUti from the BEAST package. To set the parameters, we followed the methodology as suggested by Drummond et al. (2012). The Gamma category count was set to 4. Blosum62 was used as the substitution model for the amino acid sequences and HKY with $k = 2$ for the nucleotide sequences as empirical frequencies. The Yule model was selected as the model of speciation (Harding, 1971; Nee et al., 1994). The divergence time between Zambian and European populations was set to 12,843 years, as inferred by Laurent et al. (2011). The prior for the molecular clock and speciation rates were set as gamma distributions with $\alpha = 0,001$ and $\beta = 1,000$. Further, an informative prior was set with the ZI populations as an outgroup. The mean was set to $\hat{u} = 14,000$ with a standard deviation $\sigma = 400$. The Markov Chain Monte Carlo (MCMC) options were chosen as follows: chain length 6 million, trace log(frequency) = 1,000.

Once the prior set, the phylogenetic tree was constructed with BEAST. The best tree was selected with Tree annotator from BEAST with the following options: burning percentage = 10, target tree type "Maximum clade credibility tree", node heights: "mean heights". The Bayesian parameters were estimated with Tracer 1.7 (Rambaut et al., 2018). The tree visualisation and adjustment was done with Figtree (http://tree.bio.ed.ac.uk/software/figtree/).

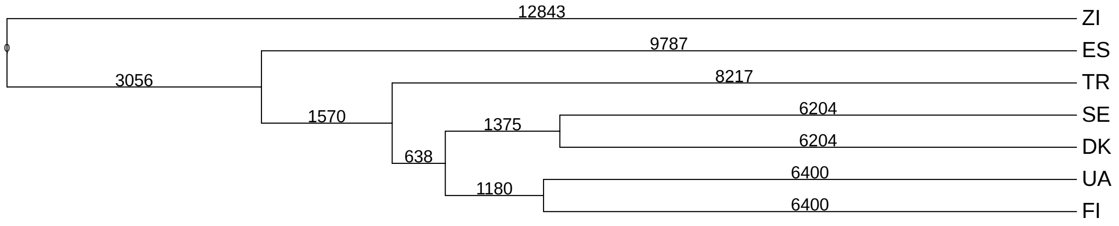


Figure S2: **Dated phylogenetic tree showing the genomic relatedness of the populations.** Branch lengths are proportional to genomic distances measured in years.

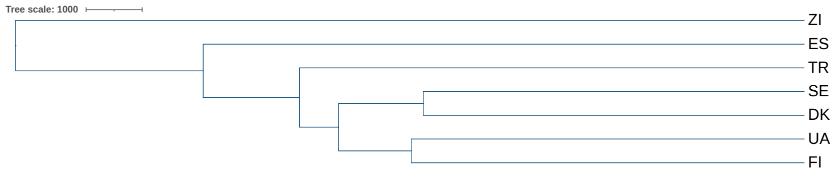


##### A.9.2 RAxML tree

The accuracy of the tree that was estimated with BEAST was tested by generating another phylogenetic tree based on the same alignment with RAxML (Stamatakis, 2016). The two estimated trees had the same topology, meaning that the inferred phylogeny of the seven populations given the data is robust.

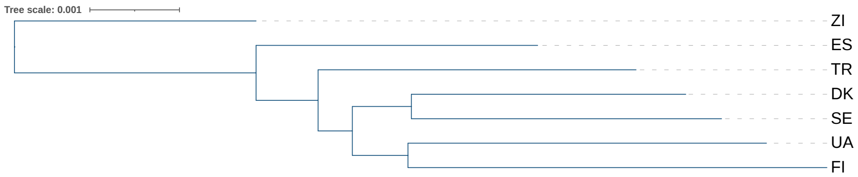


##### A.9.3 Output from BEAST

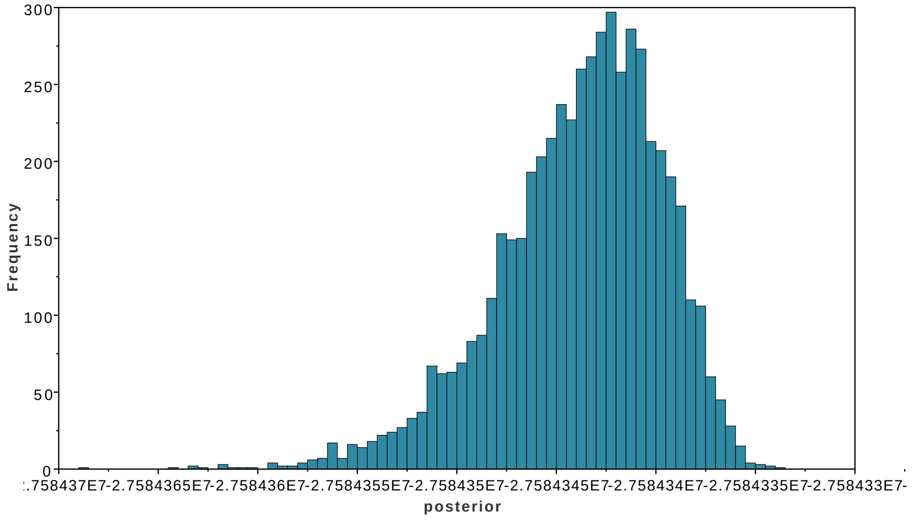

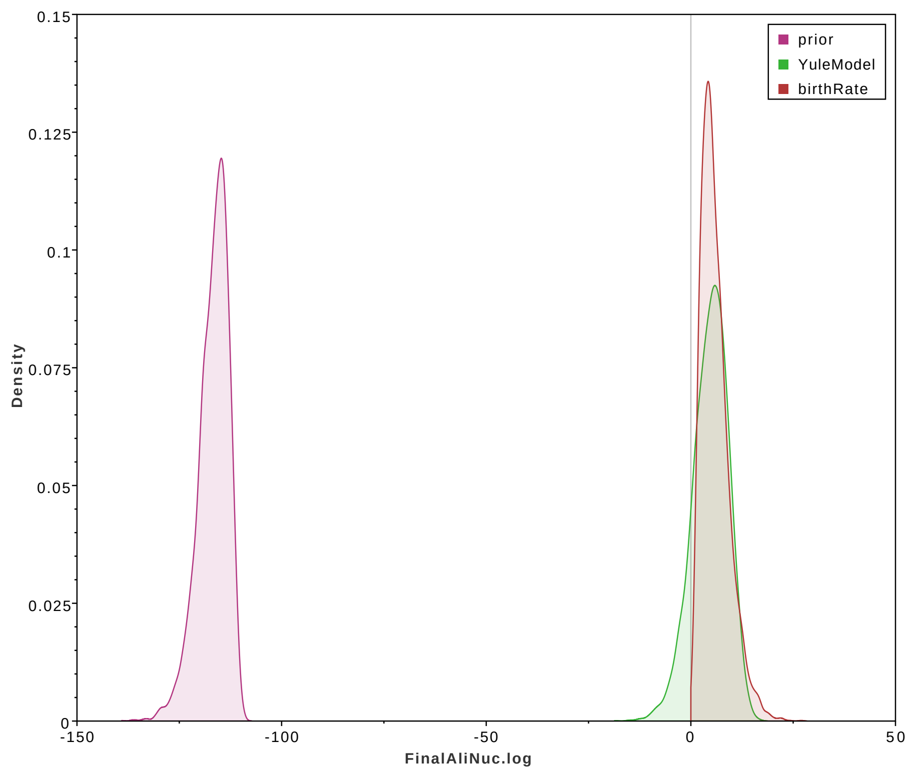

prior
YuleModel
birthRate
Density
0.15
0.125
0.1
0.075
0.05
0.025
0
-150
-100
-50
0
50
FinalAliNuc.log


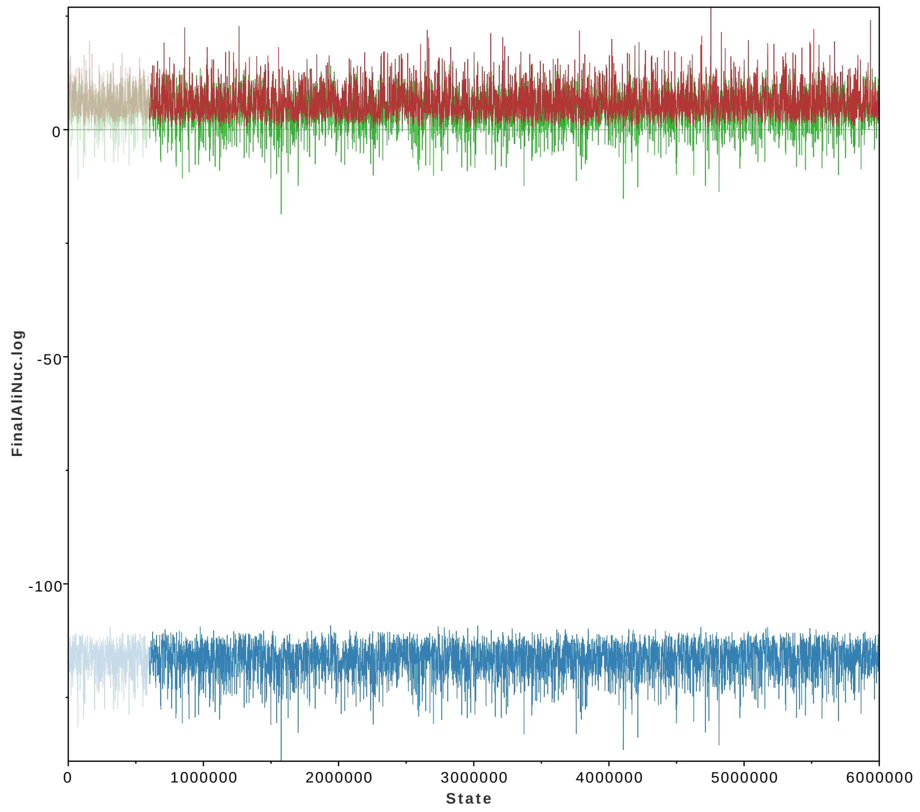

FinalAliNuc.log
0
-50
-100
0
1000000
2000000
3000000
4000000
5000000
6000000
State

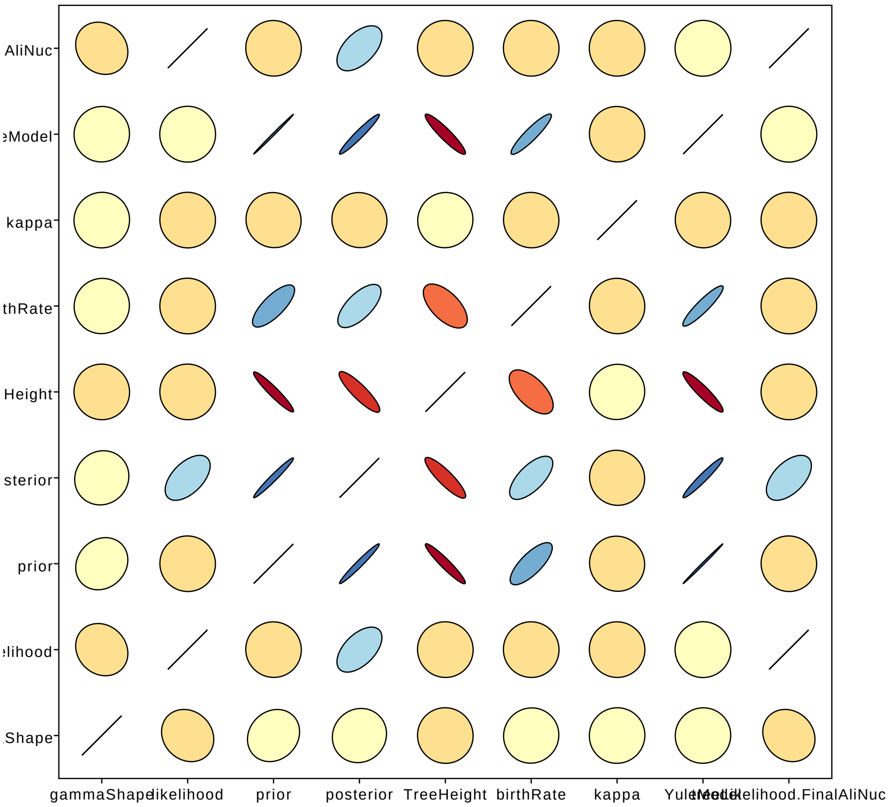

AliNuc
eModel
kappa
thRate
Height
sterior
prior
elihood
Shape
gammaShape
likelihood
prior
posterior
TreeHeight
birthRate
kappa
YuleModel
treeLikelihood.FinalAliNuc

### B Details of the parameter estimation

#### B.1 Alternative method to estimate gain and loss rates

In the main text, we have estimated the gain and loss rates based on the infinitely many genes model as originally introduced in Baumdicker et al. (2010). An adaptation of this method was proposed in Collins and Higgs (2012), where instead of the random genealogy modeled by a standard coalescent, the sample-specific dated genealogy is used to compute the frequency spectrum. To this end, a phylogeny needs to be estimated for the samples at hand, which was outlined in Section A.9. The dating of the phylogeny because of the Zambian outgroup, which can be used as the ancestral African population. Divergence times of the *D. melanogaster* out-of-Africa event have been estimated before, e.g. Laurent et al. (2011).

In this genealogy-based method, the expected transcript frequency spectrum is calculated by computing the number of transcripts gained and lost along each branch of this fixed phylogenetic tree. These gains and losses are then combined across the entire phylogeny to obtain the frequency spectrum. We follow the approach outlined in Collins and Higgs (2012). In brief, the theoretical frequency spectrum for a fixed tree with $n$ genomes (the tip nodes), and thus $n-1$ internal nodes is given by (Collins and Higgs, 2012, Eq. (14))

$$\mathbb{E}\left[G_k^n\right] = \sum_{i=1}^{2n-1} p_i(k) g(t_i), \quad \text{for } 1 \le k \le n-1, \tag{B.1}$$

where $t_i$ is the branch length leading to node $i$, $g(t_i)$ is the expected number of transcripts at node $i$ that were gained on the branch leading to it, and $p_i(k)$ is the probability that a transcript at node $i$ will be retained in $k$ tip nodes that descend from $i$. The function $g(t)$ is given by Eq. (1) in the main text and now repeated for ease of reference:

$$\frac{dg}{dt} = u - v g(t) \qquad \Rightarrow \qquad g(t) = \frac{u}{v}\left(1 - e^{-vt}\right). \tag{B.2}$$

Note that the branch leading to the root has length infinity, so that the number of transcripts in the root is given by $u/v$, the stationary value of Eq. (B.2). For the values of $p_i$, one first notes that if $i$ is a tip node, then $p_i(1) = 1$ and $p_i(k) = 0$ for $k \neq 1$. If $i$ is an internal node, one can compute the probabilities for retaining a transcript in $k$ descendants by a recursion that involves survival and loss probabilities of a transcript along the branches leading to the extant samples. The survival probability along a branch of length $t$ is given by $s(t) = e^{-vt}$ and the loss probability is $l(t) = 1 - e^{-vt}$. Denoting by $t_{i_1}$ and $t_{i_2}$ the two descending branch lengths from node $i$ that lead to nodes $i_1$ and $i_2$, the recursion is defined by (Collins and Higgs, 2012, Eqs. (16), (17))

$$\begin{aligned} p_i(k) &= s(t_{i_1}) l(t_{i_2}) p_{i_1}(k) + l(t_{i_1}) s(t_{i_2}) p_{i_2}(k) + s(t_{i_1}) s(t_{i_2}) \sum_{l=0}^{k} p_{i_1}(l) p_{i_2}(k-l), \text{ for } 1 \le k \le n; \\ p_i(0) &= \left(l(t_{i_1}) + s(t_{i_1}) p_{i_1}(0)\right)\left(l(t_{i_2}) + s(t_{i_2}) p_{i_2}(0)\right). \end{aligned} \tag{B.3}$$

As for the method that uses the standard coalescent, this procedure can be extended to multiple classes of transcripts with low and high turnover. For two classes we apply this procedure separately with the set of rates for each class and then sum up the expected frequency spectra. Moreover, if we include a class of fixed transcripts, we add the constant value $C_{\text{fixed}}$ to the value $\mathbb{E}[G_n^n]$.

In the following, we will call this estimation procedure the *fixed tree method*, and the approach based on the coalescent the *coalescent method*.

#### B.2 Initial parameter values for the parameter estimation

The optimization procedure to fit the parameters requires preset initial values. We followed the strategy suggested in Baumdicker et al. (2012) fitting the average number of observed transcripts and the observed average number of pairwise differences to the corresponding predicted theoretical values (Baumdicker et al., 2012, Eq. (1)) (ignoring the fixed number of transcripts):

$$\begin{aligned} \mathbb{E}[\text{no. of transcripts}] &= \frac{\theta_0}{\rho_0}\,, \\ \mathbb{E}[\text{no. of pairwise differences}] &= \frac{2\theta_0}{\rho_0+1}\,, \end{aligned} \tag{B.4}$$

where $\theta_0$ and $\rho_0$ denote the initial values of the gain and loss parameters during the optimization procedure. For the initial value of the fixed number of transcripts, $C_{\text{fixed}}$, we use the number of transcripts shared by all populations divided by two. In the models, where a slow and a fast class of transcripts is assumed, we set the initial values of the fast gain and loss rate according to Eq. (B.4). The initial values of the slow gain and loss rates are then obtained by dividing the fast values by 100.

In some cases the convergence procedure was not successful with these initial parameter values. We now list how we proceeded in these situations:

- all definitions: two class models (all populations and European populations only): $\theta_0 \mapsto \theta_0/4$.

#### B.3 Results from all the estimation procedures

Here, we show all the parameter estimates obtained by the different methods, fixed tree and coalescent, using different data sets arising from the different definitions, and using European samples only or all samples including the Zambian one. The results are summarized in Table B.9.

#### B.4 Results for alternative TPM threshold data sets

We conducted the same analysis as above with data sets that are generated when setting the threshold of transcription to 1 TPM or 5 TPM instead of 0.5 TPM (main text). While the gain rate is slightly reduced with these higher expression level thresholds, as is expected, the loss rate remains very similar to the one estimated with the data corresponding to a 0.5 TPM threshold. This indicates that the loss rate is independent of the transcript expression level, in line with the argument that a loss is triggered by random mutation events. The gain rate, of course, is affected by the transcript expression level in a straightforward way: the less overall transcripts we observe, the lower will the gain rate be. The parameter estimates for 1 TPM and 5 TPM data sets are presented in Tables B.10 and B.11.

| dataset | method | gain | loss | gain (slow) | loss (slow) | $C_{\text{fixed}}$ |
|---|---|---|---|---|---|---|
| **Definition 2** | | | | | | |
| all samples | 1 class – coalescent | 0.14 | $5.3\times10^{-5}$ | – | – | 71 |
| Europe | 1 class – coalescent | 0.13 | $5\times10^{-5}$ | – | – | 91 |
| all samples | 2 classes – coalescent | 0.18 | $2.1\times10^{-4}$ | 0.06 | $3.1\times10^{-5}$ | 41 |
| Europe | 2 classes – coalescent | 0.15 | $2\times10^{-4}$ | 0.06 | $3.1\times10^{-5}$ | 50 |
| all samples | 1 class – fixed tree | 0.23 | $8.9\times10^{-5}$ | – | – | 88 |
| Europe | 1 class – fixed tree | 0.23 | $9.3\times10^{-5}$ | – | – | 100 |
| all samples | 2 classes – fixed tree | 0.23 | $9.4\times10^{-5}$ | $2.1\times10^{-3}$ | $1.1\times10^{-5}$ | 0 |
| Europe | 2 classes – fixed tree | 0.23 | $9.3\times10^{-5}$ | $1.1\times10^{-4}$ | $1\times10^{-6}$ | 0 |
| **Definition 3** | | | | | | |
| all samples | 1 class – coalescent | 0.18 | $6.6\times10^{-5}$ | – | – | 75 |
| Europe | 1 class – coalescent | 0.17 | $6.3\times10^{-5}$ | – | – | 95 |
| all samples | 2 classes – coalescent | 0.34 | $3.2\times10^{-4}$ | 0.05 | $3.2\times10^{-5}$ | 50 |
| Europe | 2 classes – coalescent | 0.28 | $2.6\times10^{-4}$ | 0.05 | $3\times10^{-5}$ | 52 |
| all samples | 1 class – fixed tree | 0.26 | $9.4\times10^{-5}$ | – | – | 90 |
| Europe | 1 class – fixed tree | 0.27 | $1\times10^{-4}$ | – | – | 100 |
| all samples | 2 classes – fixed tree | 0.26 | $1\times10^{-4}$ | $2.2\times10^{-3}$ | $1.1\times10^{-5}$ | 0 |
| Europe | 2 classes – fixed tree | 0.27 | $1\times10^{-4}$ | $6.4\times10^{-4}$ | $5.1\times10^{-6}$ | 0 |

Table B.9: **Estimated *de novo* transcript gain and loss rates.** The gain and loss rates are measured as rates per year, the parameter $C_{\text{fixed}}$ is a number of transcripts. The parameters are transformed to the scale of per years, by using an effective population size of 900,000 for *D. melanogaster* in Europe (Laurent et al., 2011) and a generation time of two weeks (Fernández-Moreno et al., 2007), which gives 26 generations per year. The untransformed numerical estimates are stated in Table B.13.

| dataset | method | gain | loss | $C_{\text{fixed}}$ |
|---|---|---|---|---|
| **Definition 2** | | | | |
| all samples | 1 class – coalescent | 0.07 | $3.5\times10^{-5}$ | 42 |
| Europe | 1 class – coalescent | 0.07 | $3.3\times10^{-5}$ | 50 |
| **Definition 3** | | | | |
| all samples | 1 class – coalescent | 0.12 | $6.1\times10^{-5}$ | 57 |
| Europe | 1 class – coalescent | 0.11 | $5.9\times10^{-5}$ | 74 |

Table B.10: **Estimated *de novo* transcript gain and loss rates for the 1 TPM threshold data set.** The gain and loss rates are measured as rates per year, the parameter $C_{\text{fixed}}$ is a number of transcripts. The parameters are transformed to the scale of per years, by using an effective population size of 900,000 for *D. melanogaster* in Europe (Laurent et al., 2011) and a generation time of two weeks (Fernández-Moreno et al., 2007), which gives 26 generations per year.

| dataset | method | gain | loss | $C_{\text{fixed}}$ |
|---|---|---|---|---|
| **Definition 2** | | | | |
| all samples | 1 class – coalescent | 0.04 | $4.1\times10^{-5}$ | 19 |
| Europe | 1 class – coalescent | 0.03 | $4\times10^{-5}$ | 29 |
| **Definition 3** | | | | |
| all samples | 1 class – coalescent | 0.05 | $5.5\times10^{-5}$ | 21 |
| Europe | 1 class – coalescent | 0.04 | $5.3\times10^{-5}$ | 29 |

Table B.11: **Estimated *de novo* transcript gain and loss rates for the 5 TPM threshold data set.** The gain and loss rates are measured as rates per year, the parameter $C_{\text{fixed}}$ is a number of transcripts. The parameters are transformed as explained in the caption of Table B.10.

#### B.5 Normalization of gain rates per genomic region

To compare the transcript gain rates between different genomic regions, we need to normalize the estimated gain rates. The reason is that regions with a high coverage in the genome, e.g. intergenic regions, have a larger chance to gain a *de novo* transcript due to random mutations than regions with a low coverage, e.g. non-coding RNA regions. We therefore normalize the gain rates, denoted by $u_{\text{norm}}$, to reflect the gain of a transcript per genome coverage. The coverages of the different genomic regions are as follows: exon longer = 15%, intergenic = 64%, intronic = 23%, non-coding RNA = 9%, antisense = 15%. Note that we exclude the pseudogenic regions from this analysis because of their overall low number of transcripts and extremely long transcript lengths compared to the other genomic regions.

The regions of potential gain for intergenic and intronic regions are calculated by their relative coverage of the entire genome. The other regions are more complicated to compute because transcripts of these classes overlap an intergenic region and their classification, e.g. a transcript classified as 'exon longer' overlaps with an intergenic region and a gene in the direction of its transcription. We therefore apply the following reasoning to estimate the relative genomic coverage of these overlapping regions. We first compute the median transcript length, which is 1,467 base pairs (bp). We then multiply this number with the number of occurrences of this genomic region. Based on the *D. melanogaster* reference genome, there are 13,963 annotated genes in *D. melanogaster* and 4,015 non-coding RNAs. Among the genes, 16% are estimated to overlap, reducing the number to 11,729 (Lee and Chang, 2013). In addition, we multiply the number of genes or non-coding RNAs by two because a transcript can overlap on both sides. This value is then divided by the total genome length of *D. melanogaster*, which is 275,134,968 bp (forward and backward read of the chromosome length). For example, to compute the coverage of 'antisense' we have

$$\frac{\overbrace{1,467}^{\text{median length of a transcript}} \times \overbrace{11,729}^{\text{no. of genes}} \times \overbrace{2}^{\text{overlap with left or right boundary of the gene}}}{275,134,968} = 0.15\,. \tag{B.5}$$

The normalized gain rate is then given by $u_{\text{norm}} = u/\text{coverage}$.

For comparison with the results from the coalescent-based method, we also show the position-specific parameter estimates from the fixed tree method in Table B.12.

| genomic region | **Definition 2** $u$ | $u_{\text{norm}}$ | $v$ | $C_{\text{fixed}}$ | **Definition 3** $u$ | $u_{\text{norm}}$ | $v$ | $C_{\text{fixed}}$ |
|---|---|---|---|---|---|---|---|---|
| exon longer | 0.03 | 0.19 | $9.8\times10^{-5}$ | 27 | 0.04 | 0.29 | $10^{-4}$ | 30 |
| intergenic | 0.06 | 0.1 | $1.1\times10^{-4}$ | 7 | 0.08 | 0.12 | $1.2\times10^{-4}$ | 7 |
| intronic | 0.01 | 0.04 | $1.2\times10^{-4}$ | 0 | 0.02 | 0.1 | $1.2\times10^{-4}$ | 3 |
| non-coding RNA | 0.01 | 0.13 | $1.1\times10^{-4}$ | 6 | 0.03 | 0.29 | $1.1\times10^{-4}$ | 8 |
| antisense | 0.16 | 1.1 | $1.1\times10^{-4}$ | 29 | 0.19 | 1.25 | $1.1\times10^{-4}$ | 22 |

Table B.12: **Estimated *de novo* transcript gain and loss rates per genomic region.** We have used the one class, fixed-tree method to estimate the gain and loss rates of transcripts on the European samples. Parameter estimates of $u$ and $v$ are per year. To compare the gain rates between different genomic regions, we have normalized the gain rates according to the coverage of the region in the genome. Normalization is done by rescaling the gain rates as described in the text, i.e. $u_{\text{norm}} = u/\text{coverage}$.

#### B.6 Untransformed parameter estimates

In the main text we transformed the parameters to rates per year (Table 3 in the main text). Here, we state the non-transformed parameters as obtained from the numerical estimation procedure (Table B.13).

| dataset | method | $\theta_f$ or $u_f$ | $\rho_f$ or $v_f$ | $\theta_s$ or $u_s$ | $\rho_s$ or $v_s$ | $C_{\text{fixed}}$ |
|---|---|---|---|---|---|---|
| **Definition 1** | | | | | | |
| all populations | 1 class – coalescent | 6,283 | 2.31 | – | – | 61 |
| Europe | 1 class – coalescent | 5,777 | 2.16 | – | – | 65 |
| all populations | 2 classes – coalescent | 6,290 | 2.5 | 1560 | 0.58 | 0 |
| Europe | 2 classes – coalescent | 5,794 | 2.25 | 63 | 0.37 | 0 |
| all populations | 1 class – fixed tree | 2,764 | 1.04 | – | – | 115 |
| Europe | 1 class – fixed tree | 2,125 | 0.81 | – | – | 127 |
| all populations | 2 classes – fixed tree | 2,774 | 1.1 | 34 | 0.13 | 0 |
| Europe | 2 classes – fixed tree | 2,124 | 0.82 | 2.43 | 0.02 | 0 |
| **Definition 2** | | | | | | |
| all populations | 1 class – coalescent | 9,597 | 3.69 | – | – | 71 |
| Europe | 1 class – coalescent | 8,824 | 3.48 | – | – | 91 |
| all populations | 2 classes – coalescent | 12,178 | 15 | 3,844 | 2.15 | 41 |
| Europe | 2 classes – coalescent | 10,550 | 14 | 3,893 | 2.13 | 50 |
| all populations | 1 class – fixed tree | 2,947 | 1.14 | – | – | 88 |
| Europe | 1 class – fixed tree | 2,297 | 0.91 | – | – | 100 |
| all populations | 2 classes – fixed tree | 2,981 | 1.21 | 27 | 0.14 | 0 |
| Europe | 2 classes – fixed tree | 2,298 | 0.91 | 1.13 | 0.01 | 0 |
| **Definition 3** | | | | | | |
| all populations | 1 class – coalescent | 12,504 | 4.57 | – | – | 75 |
| Europe | 1 class – coalescent | 11,575 | 4.34 | – | – | 95 |
| all populations | 2 classes – coalescent | 23,783 | 22 | 3,676 | 2.22 | 50 |
| Europe | 2 classes – coalescent | 19,609 | 18 | 3,326 | 2.08 | 52 |
| all populations | 1 class – fixed tree | 3,284 | 1.21 | – | – | 90 |
| Europe | 1 class – fixed tree | 2,601 | 0.98 | – | – | 100 |
| all populations | 2 classes – fixed tree | 3,342 | 1.28 | 28 | 0.14 | 0 |
| Europe | 2 classes – fixed tree | 2,609 | 0.99 | 6.3 | 0.05 | 0 |

Table B.13: **Estimated *de novo* transcript gain and loss rates.** Coalescent parameter estimates are per $2N_e$ generations. Parameter estimates for the fixed tree are per 12,843 years when using all populations and per 9,787 years when using only the European populations.
